# Comparative analysis of gonadal hormone receptor expression in the house mouse, meadow vole, and prairie vole brain

**DOI:** 10.1101/2023.05.12.540606

**Authors:** Katherine A. Denney, Melody V. Wu, Simón(e) D. Sun, Soyoun Moon, Jessica Tollkuhn

## Abstract

The socially monogamous prairie vole (*Microtus ochrogaster*) and promiscuous meadow vole (*Microtus pennsylvanicus*) are closely related, but only prairie voles display long-lasting pair bonds, biparental care, and selective aggression towards unfamiliar individuals after pair bonding. These social behaviors in mammals are largely mediated by steroid hormone signaling in the social behavior network (SBN) of the brain. Hormone receptors are reproducible markers of sex differences that can provide more information than anatomy alone, and can even be at odds with anatomical dimorphisms. We reasoned that behaviors associated with social monogamy in prairie voles may emerge in part from unique expression patterns of steroid hormone receptors in this species, and that these expression patterns would be more similar across males and females in prairie than in meadow voles or the laboratory mouse. To obtain insight into steroid hormone signaling in the developing prairie vole brain, we assessed expression of estrogen receptor alpha (*Esr1*), estrogen receptor beta (*Esr2*), and androgen receptor (*Ar*) within the SBN, using *in situ* hybridization at postnatal day 14 in mice, meadow, and prairie voles. We found species-specific patterns of hormone receptor expression in the hippocampus and ventromedial hypothalamus, as well as species differences in the sex bias of these markers in the principal nucleus of the bed nucleus of the stria terminalis. These findings suggest the observed differences in gonadal hormone receptor expression may underlie species differences in the display of social behaviors.

## Introduction

Mammalian species show a wide diversity of social behaviors. This natural variation can be leveraged to identify the unique circuitry that imparts a behavior of interest. Several vole species are established model organisms for the study of sociality (He et al., 2019; Insel and Young, 2001; Lee and Beery, 2019; Sadino and Donaldson, 2018; Carter et al., 1995). There are stark behavioral differences between monogamous and promiscuous vole species: prairie voles display social monogamy, biparental care, and form long-lasting pair bonds, which rarely occur in mammals, while promiscuous meadow voles are socially non-monogamous and uniparental. This contrast provides a framework for the study of the neural circuitry that enables social attachment behaviors. Extensive comparative studies between prairie voles and non-monogamous vole species demonstrate that the unique distribution of oxytocin receptor (*Oxtr*) and vasopressin receptor (*Avpr1a*) in prairie voles underlies pair bond formation (Insel et al., 1991; Lim et al., 2004; Ophir et al., 2012; Wang et al., 1996; Williams et al., 1994; Winslow et al., 1993; Young et al., 2011). However, how the prairie vole brain develops the capacity to form pair bonds remains an open question: what events instruct *Oxtr* and *Avpr1a* expression and what other genes might show distinct patterns of gene expression in prairie voles and other monogamous species?

Social monogamy is accompanied by decreased behavioral sex differences in prairie voles compared to mice and rats. Male prairie voles care for offspring and juveniles and show high levels of spontaneous alloparenting (Bales et al., 2006; Carter and Getz, 1993; Getz et al., 1981; Kramer et al., 2009; Lonstein and De Vries, 2000). Prairie voles also display similar levels of aggression in females and males. Males attack novel conspecifics at low levels, but upon pairbond formation, both sexes show selective aggression towards strangers (Getz et al., 1981; Lee and Beery, 2022; Tickerhoof et al., 2020; Wang et al., 1997; Young et al., 2011). Thus, a pair-bond encompasses both prosocial (towards the partner) and antisocial (towards novel conspecifics) behaviors.

Social behaviors such as aggression, mounting, lordosis, and parenting behaviors, are mediated by the social behavior network (SBN), which processes pheromonal cues to determine social context and select the appropriate response (Chen and Hong, 2018; Goodson, 2005; Newman, 1999; Wallace et al., 2023). While females and males typically display sex differences in the frequency and intensity of these outputs, the ability to engage in most social behaviors is present in both sexes (Wei et al., 2021). For example, both female and male mice show aggression towards intruders, but male mice attack only male intruders, while female mice attack both sexes, but only while lactating (Hashikawa et al., 2018; Liu et al., 2021; Lonstein and Gammie, 2002; St. John and Corning, 1973). Similarly, although females are the primary caregivers in the majority of mammalian species, fathers can express many of the same stereotyped parental behaviors as mothers particularly in monogamous species (Bales and Saltzman, 2016; Dewsbury, 1985; Oliveras and Novak, 1986; Zilkha et al., 2017). Therefore, the circuitry of innate social behaviors must be shared between sexes yet facilitate sex-differential responses to social context.

In many species, including humans, several regions of the SBN show sexual dimorphism in cell density, cell number, volume, projection pattern, or gene expression, particularly the sexually dimorphic nucleus of the preoptic area (SDN-POA), the principal nucleus of the bed nucleus of the stria terminalis (BNSTpr), and anteroventral periventricular nucleus (AVPV) (Allen and Gorski, 1990; Gorski et al., 1978; Kelly et al., 2013; Simerly et al., 1985; Tsukahara and Morishita, 2020). In mice and rats, these dimorphisms are organized by perinatal testosterone signaling in males. Circulating testosterone is locally converted to 17β-estradiol in select neuronal populations that express aromatase and this neural estradiol drives brain sexual differentiation (Balthazart and Ball, 1998; Juntti et al., 2010; Lephart, 1996; MacLusky and Naftolin, 1981; Naftolin and Ryan, 1975; Wu et al., 2009). Intriguingly, monogamous species such as prairie voles also show decreased sexual dimorphism in the brain, as well as in anatomical measures such as body weight and anogenital distance (Campi et al., 2013; Dewsbury et al., 1980; Heske and Ostfeld, 1990; Shapiro et al., 1991). In addition, prairie vole social behaviors are resistant to early life testosterone manipulations, as if the classic organization and activation model is operating under different constraints in this species. Like other rodents, prairie vole males undergo a perinatal testosterone surge (Lansing et al., 2013), but in contrast to rats, neonatal orchiectomy does not abolish adult male sexual behavior. Instead, postnatal testosterone *reduces* androgen induced mounting behavior in adult males (Roberts et al., 1997). Perinatal testosterone also does not masculinize the expression of arginine vasopressin (*Avp*) in the BNST and medial amygdala (MeA) (Lonstein et al., 2002). These findings suggest that prairie vole brain sexual differentiation follows a distinct trajectory compared to both mice and rats.

Behaviors associated with social monogamy may emerge in part from unique expression patterns of gonadal steroid hormone receptors in prairie voles. Hormone receptors are reproducible markers of sex differences that add additional information compared to anatomy alone, and can even be at odds with anatomical dimorphisms. In adult mice and rats, *Esr1* and progesterone receptor (*Pgr*) are more abundant in females, even within the BNSTpr and medial preoptic area (MPOA), which contain more neurons in males (Gegenhuber et al., 2022; Quadros and Wagner, 2008; Xu et al., 2012; Yang et al., 2013; Yokosuka et al., 1997). Furthermore, in the BNST, *Esr1*-expressing neurons are more transcriptionally different between sexes than neurons not expressing hormone receptors (Gegenhuber et al., 2022). Variation in steroid hormone signaling has previously been implicated as a driver of social behavior evolution (Adkins-Regan, 2012; Hoke et al., 2019; Kelly and Vitousek, 2017; Young and Crews, 1995), but there have been few comparative studies of qualitative differences in hormone receptor expression between rodent species (Cushing, 2016; Cushing et al., 2004; Cushing and Wynne-Edwards, 2006). Moreover, although sex differences in ERα immunostaining in adult prairie voles have been reported previously (Cushing et al., 2004; Hnatczuk et al., 1994), expression of *Ar* and *Esr2* has not been previously investigated.

To obtain insight into prairie vole brain sexual differentiation, we assessed expression of *Esr1, Esr2*, and *Ar* within the SBN, using in situ hybridization (ISH) at postnatal day 14 (P14) in mice, prairie, and meadow voles. The inclusion of meadow voles permits comparisons within two *Microtus* species and provides a behavioral intermediate between polygamous mice and monogamous prairie voles: meadow voles show a partner preference for same-sex peers and engage in seasonal social group living (Anacker et al., 2016; Beery, 2019; Beery et al., 2014; Lee and Beery, 2022). We selected ISH to allow a consistent methodology across all three genes. Immunostaining for ERβ has been historically complicated by antibody variability; a previous comprehensive characterization of ERβ expression in mice utilized a transgenic *Esr2*-GFP line (Andersson et al., 2017; Nelson et al., 2017; Zuloaga et al., 2014). We chose P14 as this time point is beyond the closure of the postnatal sensitive period for brain sexual differentiation in mice and rats (MacLusky and Naftolin, 1981; McCarthy, 2008; Simerly, 2002), precedes pubertal hormone cycles, and coincides with critical periods for social learning in rodents (Hammock, 2014; Newmaster et al., 2020). We additionally investigated sex differences in hormone receptor expression to determine if prairie voles show decreased sexual dimorphism compared to mice.

## Results

### Species and Sex Differences in *Esr1* Expression

Although overall localization of *Esr1* transcripts within the SBN was largely similar across species, several differences were readily apparent in the regions schematized in Figure 1A. We first observed minimal *Esr1* staining within the mouse hippocampus (Figure 1B,C). To corroborate this low signal, we turned to a large snRNA-seq dataset generated by the Allen Institute that encompasses the entire mouse isocortex and hippocampal formation (Yao et al., 2021). Across 47 clusters of hippocampal neurons, expression of *Esr1* appears in only two ventral CA3 clusters, and a single CA1 cluster, with relatively low expression even within these clusters (Figure 1D).

**Figure 1.**
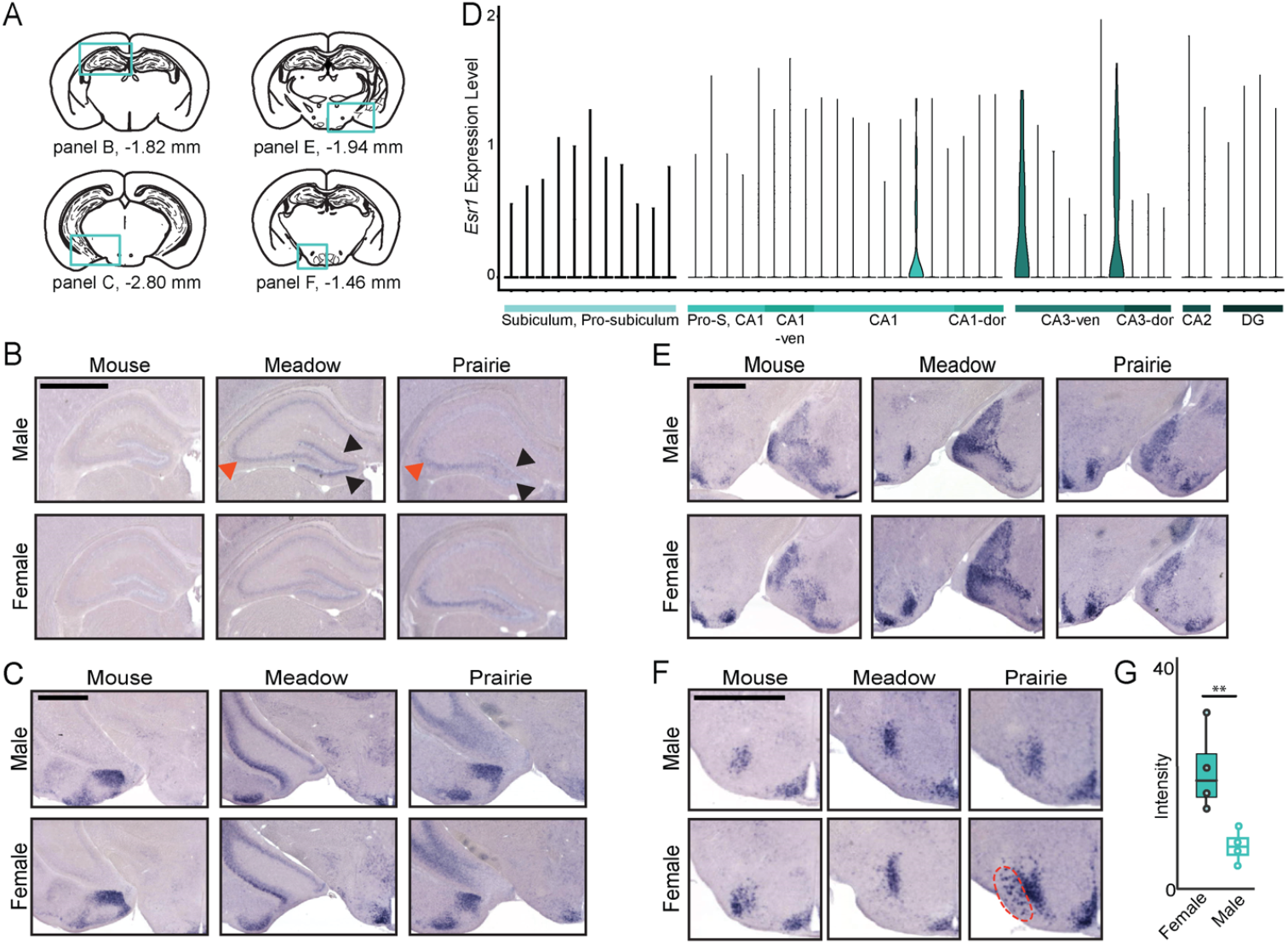
Species differences in *Esr1* expression. **(A)** Schematics of mouse coronal sections for dorsal hippocampus (dHPC), ventral hippocampus (vHPC), medial amygdala (MeA), and ventromedial hypothalamus (VMH) (Paxinos and Franklin, 2019). **(B-C)** *In situ* hybridization (ISH) for *Esr1* expression in dHPC (B) and vHPC (C). Red arrowheads denote CA3, black arrowheads denote dentate gyrus. Note species differences in vHPC. Male (top panels) and female (bottom panels) are shown for mouse, meadow vole, and prairie vole (left to right). **(D)** Violin plot of *Esr1* expression across hippocampal transcriptional clusters organized by anatomical subregion. See Supplementary Table 2 for subclass abbreviations and cluster identity. **(E-F)** ISH for *Esr1* expression in MeA (E) and VMH (F). Red dotted lines outline VMHvll. **(G)** Quantification of VMHvll staining intensity in prairie vole females and males. Boxplot denotes median and 1st & 3rd quartiles, ** p<0.01. Scale bars = 1000 μm.

In contrast, *Esr1* signal was strong in the hippocampus of both meadow and prairie voles, with striking enrichment in distinct subregions. In the dorsal hippocampus of meadow, but not prairie voles, *Esr1* is particularly abundant in the dentate gyrus (black arrowheads), yet is strongly expressed within CA3 only in prairie voles (red arrowheads) (Figure 1B). In the ventral hippocampus, there is staining in the amygdalo-hippocampal and amydalopiriform area in both meadow and prairie vole (Figure 1C). The full range of staining intensity is shown on full sections in Supplementary Figure 1A. We noted an anatomical difference in that the ventral portion of the hippocampus extends dorsally from the ventral surface moving through posterior coronal sections (Supplementary Figure 1B) in meadow voles. Cortical staining of *Esr1* was minimal in all three species and is not shown by ISH; transcriptomic taxonomy neighborhood-specific expression in mouse by snRNAseq is shown for inhibitory cortical cell types (Supplementary Figure 1C) and excitatory cortical cell types (Supplementary Figure 1D-H).

*Esr1* is expressed in the medial amygdala (MeA), ventromedial hypothalamus ventrolateral region (VMHvl), and arcuate nucleus as shown in Figure 1E. We found an unexpected sex difference in the localization of *Esr1* transcripts in prairie voles. In addition to the intense VMHvl staining typically seen in mice, females also express *Esr1* in a more lateral population (Figure 1F, red dashed line). Quantification of *Esr1* expression in this VMHvll region showed a significant sex difference in a log ratio test (95% CI (−0.724, -0.200); p=0.0014; N=4 animals per sex, Figure 1G, Supplementary Figure 2A). This region was quantified only in prairie voles because the anatomic VMHvll subdivision was not sufficiently distinguishable from the principal VMHvl expression in mice or meadow voles, suggesting a novel, prairie vole-specific sexual dimorphism.

Within the SBN of mice and rats, *Esr1* levels are higher in females compared to males, particularly within the VMHvl, POA, and BNSTpr (Gegenhuber et al., 2022; Ikeda et al., 2003; Kelly et al., 2013; Xu et al., 2012). While assessing species differences in *Esr1* expression, we noted that sex differences appeared more subtle in both species of voles compared to mice. We selected the BNSTpr for quantitation as sex differences in this region have not previously been investigated in voles (Campi et al., 2013; Shapiro et al., 1991). As expected, we observed a robust sex difference in *Esr1* levels in the BNSTpr of mice (95% CI (−0.203,-0.068); p=0.00054, N=5 animals per sex) (Figure 2). However, we did not detect a significant difference between sexes in either meadow (95% CI (−0.039,0.088); p=0.32; N=5 animals per sex) or prairie voles (95% CI (−0.359,0.224); p=0.49; N=4 animals per sex). The log ratio of male to female expression was not significantly different from zero in either vole species (Figure 2E). The quantified region (5 consecutive sections of principal nucleus only) is outlined in Supplementary Figure 2B.

**Figure 2.**
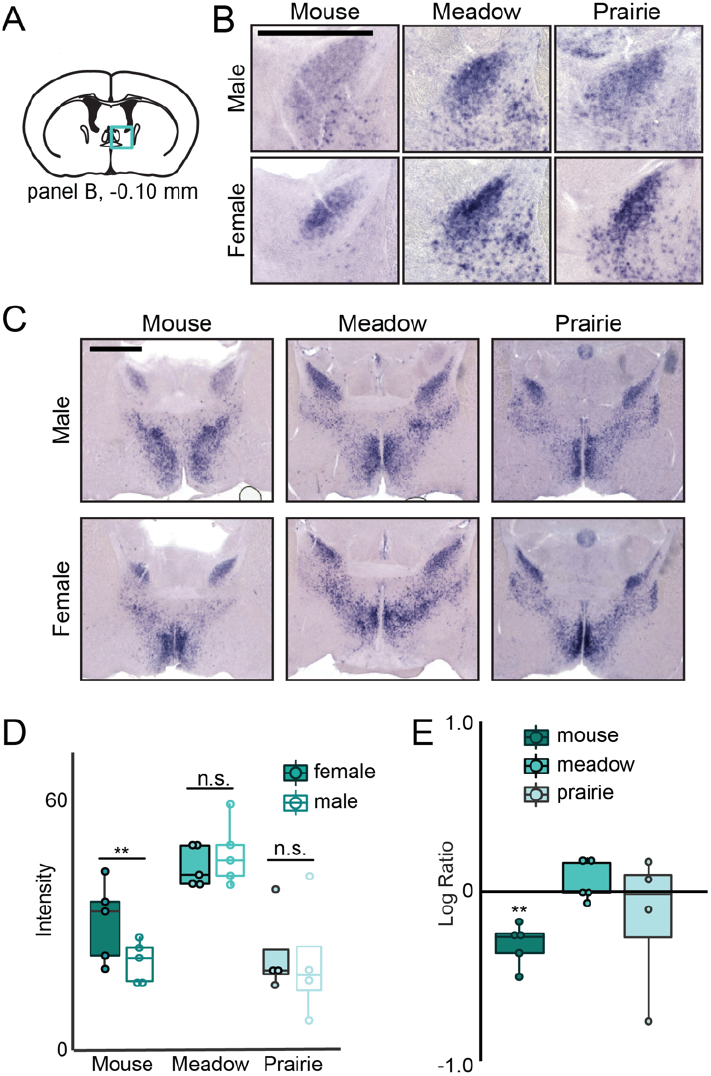
Sex differences in *Esr1* expression. **(A)** Schematic of coronal section for bed nucleus of stria terminalis prin-cipal nucleus (BNSTpr) *Esr1* expression and quantification. **(B)** ISH for *Esr1* in BNSTpr. Male (top panels) and female (bottom panels) are shown for mouse, meadow vole, and prairie vole (left to right). **(C)** ISH for *Esr1* in BNST and preoptic area (POA) of the hypothalamus. **(D)** Quantification of BNSTpr *Esr1* expression in males and females of each species (N=5, mouse, meadow vole; N=4, prairie vole). Boxplot denotes median and 1st & 3rd quartiles, ** p<0.01. **(E)** Within-batch male:female log ratios of *Esr1* expression in BNSTpr. ** denotes significant difference from log ratio of zero. Scale bars = 1000 μm

### Species and Sex Differences in *Ar* Expression

In mice and rats, *Ar* expression is more widespread than that of *Esr1* (Brock et al., 2015; Simerly et al., 1990). We performed ISH for *Ar* in our three species and identified differences within the regions schematized in Figure 3A. As with *Esr1*, we found pronounced species differences in hippocampal *Ar* expression (Figure 3B). While *Ar* is strongly expressed in the CA1 of all three species, high expression within CA2 is only seen in mice (gray arrows). In the CA3 (red arrows) and dentate gyrus (black arrows), *Ar* expression mirrors that of *Esr1*, with more expression in prairie voles compared to other species in CA3 and in dentate gyrus of meadow voles, as well as mice, compared to prairie voles. The strong expression of *Ar* within the mouse hippocampus is also seen by snRNA-seq, where *Ar* is present in most excitatory cell types (Figure 3C).

**Figure 3.**
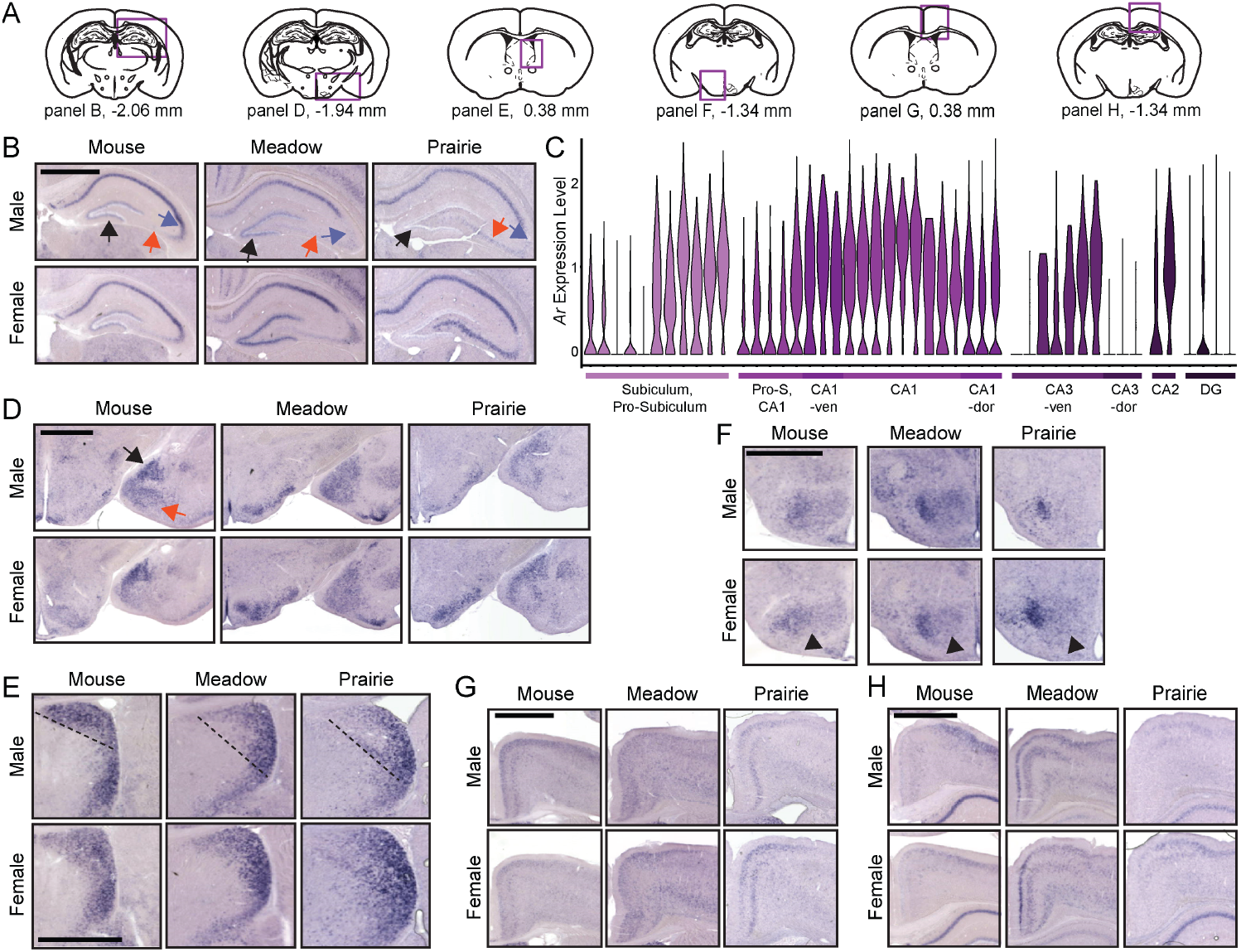
Species differences in *Ar* expression. **(A)** Schematics of coronal sections for dorsal hippocampus (dHPC), medial amygdala (MeA), lateral septum (LS), ventromedial hypothalamus (VMH), anterior cingulate cortex (ACC), and retrosplenial area. **(B)** ISH for *Ar* in dHPC. Black arrows denote dentate gyrus, red arrows denote CA3, and blue-gray arrows denote CA2. Male (top panels) and female (bottom panels) are shown for mouse, meadow vole, and prairie vole (left to right). **(C)** Violin plots of *Ar* expression in hippocampal transcriptional cell type clusters organized by subregion. See Supplementary Table 2 for subclass abbreviations and cluster identity. **(D-H)** ISH for *Ar* in MeA **(D)**, LS **(E)**, VMH **(F)**, ACC **(G)**, and retrosplenial area **(H)**. Black arrow indicates MeApd, red arrow indicates MeApv. Dashed line indicates the dorsal LS region of interest. Black arrowheads indicate arcuate nucleus. Scale bars=1000 μm.

Generally, *Ar* appears more diffusely expressed in prairie voles than meadow voles or mice, particularly in the MeA (Figure 3D). The full range of staining intensity can be seen in Supplementary Figure 3A. However, the density of subregions within the MeA is more defined in mice than in either vole species, with a clear distinction between the posterodorsal nucleus (MeApd, black arrow) and posteroventral nucleus (MeApv, red arrow). The distribution of *Ar* expression in the lateral septum (LS) is different between mouse and vole species, with *Ar* expression taking up a broader area of dorsal LS in voles than in mice (Figure 3E). In mice, *Ar* expression is restricted to a thinner region of the dorsal lateral septum, while in voles the *Ar* expression in the dorsal region protrudes more medially and ventrally.

In the ventromedial hypothalamus, *Ar* is expressed throughout the VMH (including dorsomedial and central VMH) in the mouse (Figure 3F, left). In the prairie vole, expression is more restricted to the VMHvl (Figure 3F, right). Meadow voles have an intermediate expression pattern with broad expression throughout the VMH and enrichment of expression in VMHvl (Figure 3F, center). Additionally, although *Ar* is expressed in the arcuate nucleus in mice, it is largely absent from meadow and prairie voles (Figure 3F, black arrowheads).

*Ar* is much more prevalent throughout the cortex compared to *Esr1* (Supplementary Figure 3B-D), both in the anterior cingulate cortex (Figure 3G), as well as the more posterior retrosplenial area (Figure 3H), Accordingly, in snRNA-seq data, *Ar* is present within multiple inhibitory neuronal types, barring parvalbumin interneurons (Supplementary Figure 3C), and is also prevalent in excitatory populations, particularly the deep layers (Supplementary Figure 3E-H).

Both mice and meadow voles had a pronounced difference in *Ar* expression in the BNSTpr with male expression higher than females (mouse 95% CI (−0.003,0.237); p=0.027, meadow 95% CI (0.033,0.183); p=0.0041), while prairie voles had no significant differences in BNSTpr *Ar* expression (95% CI (−0.225,0.231); p=0.97), (Figure 4, N=5 animals per sex for all species). The region of BNSTpr quantification is outlined in 5 consecutive BNSTpr sections in Supplementary Figure 4.

**Figure 4.**
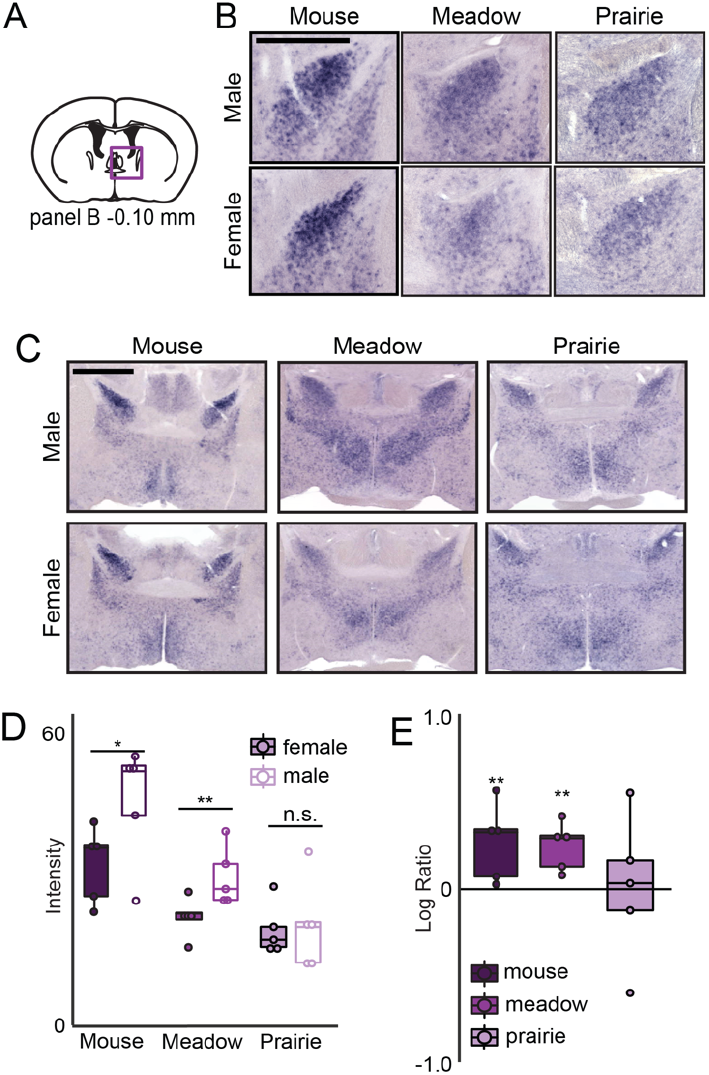
Sex differences in *Ar* expression. **(A)** Schematic of coronal section for BNSTpr *Ar* expression and quantifi-cation. **(B)** ISH for *Ar* in BNSTpr. Male (top panels) and female (bottom panels) are shown for mouse, meadow vole, and prairie vole (left to right). **(C)** ISH for *Ar* in BNST and preoptic area (POA) of the hypothalamus. **(D)** Quantification of BNSTpr *Ar* expression in males and females of each species (N=5). Boxplot denotes median and 1st & 3rd quartiles, * p<0.05, ** p<0.01. **(E)** Within-batch male:female log ratios of *Ar* expression in BNSTpr. ** denotes significant difference from log ratio of zero. Scale bars = 1000 μm.

### Species and Sex Differences in *Esr2* Expression

*Esr2* expression in all three species is extremely sparse within the span of sections that we assessed (Figure 5A) as well as by snRNAseq in the mouse hippocampus (Figure 5B). *Esr2* is highly expressed in the paraventricular nucleus of the hypothalamus (PVN) of all three species (Figure 5C). Notably, PVN expression in meadow voles extends through nearly twice as many coronal sections as in mouse and prairie vole, indicating a larger anterior-posterior distribution of *Esr2* expression in meadow vole PVN (not shown). *Esr2* is expressed in MeApd in mice, and more broadly throughout the posterior MeA in both vole species (Figure 5D). *Esr2* is sparsely expressed in the AVPV (Figure 5E). However, it appears to be expressed more densely in female meadow vole AVPV compared to all other species and sexes. This relative expression pattern is consistent throughout the anterior-posterior extent of the AVPV. We did not detect hippocampal *Esr2* expression in any species (Figure 5F), shown on the same sections as MeA staining in Supplementary Figure 5A. In the cortex, *Esr2* is similarly sparsely expressed, with negligible expression in all inhibitory cell types (Supplementary Figure 5B) and most excitatory cell types (Supplementary Figures 5C-G), with low expression in only two excitatory clusters of layer 2/3 IT PPP and IT ENTm.

**Figure 5.**
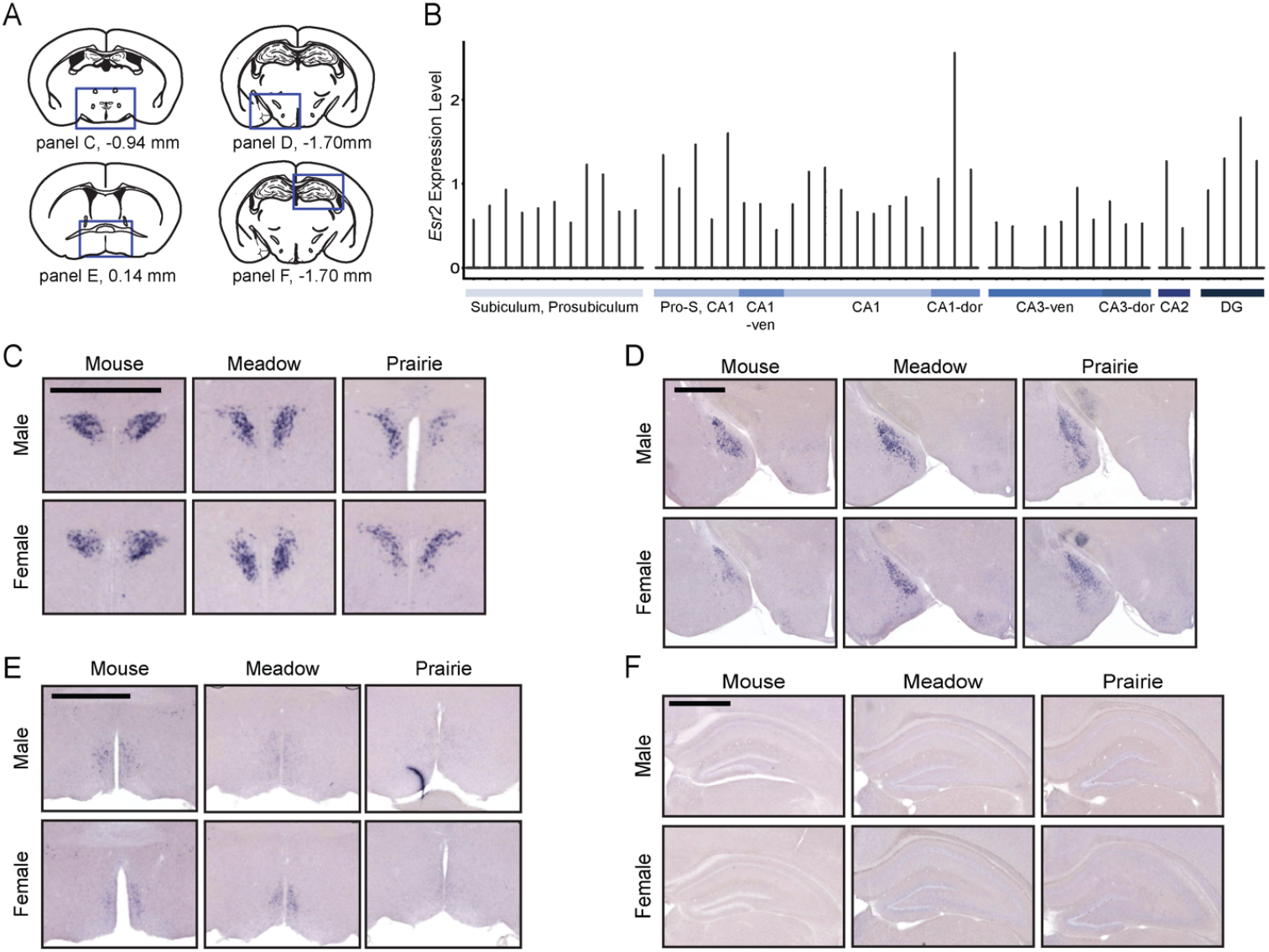
Species differences in *Esr2* expression. **(A)** Schematics of coronal sections for the paraventricular nucleus of the hypothalamus (PVN), medial amygdala (MeA), anteroventral periventricular nucleus (AVPV), and dorsal hippocampus (dHPC). **(B)** Violin plots of *Esr2* expression across hippocampal transcriptional cell type clusters organized by anatomical subregion. See Supplementary Table 2 for subclass abbreviations and cluster identity. **(C-F)** ISH for *Esr2* in PVN **(C)**, MeA **(D)**, AVPV **(E)**, and dHPC **(F)**. Male (top panels) and female (bottom panels) are shown for mouse, meadow vole, and prairie vole (left to right). Scale bars = 1000 μm.

In the BNSTpr, we expected female *Esr2* expression to be higher than male expression, based on previous BNSTpr snRNA-seq in mouse pups (Gegenhuber et al., 2022). This pattern held true for mice (95% CI (−0.077, -0.011); p=0.0052; Figure 6, N=4 animals per sex), but we saw higher expression of *Esr2* in male meadow voles than female meadow voles (95% CI (0.031,0.172); p=0.0037; Figure 6, N=4 animals per sex). Prairie voles show no significant expression differences (95% CI (−0.201,0.245); p= 0.77; Figure 6, N=4 animals per sex) between sexes in BNSTpr. In mice and prairie voles, we noted a distinctive “hole” in *Esr2* signal in males, consistent with previously described expression of an Esr2-Cre line (Zhou et al., 2023). The region of BNSTpr quantification is outlined in 5 consecutive BNST sections in Supplementary Figure 6. described expression of an Esr2-Cre line (Zhou et al., 2023). The region of BNSTpr quantification is outlined in 5 consecutive BNST sections in Supplementary Figure 6.

**Figure 6.**
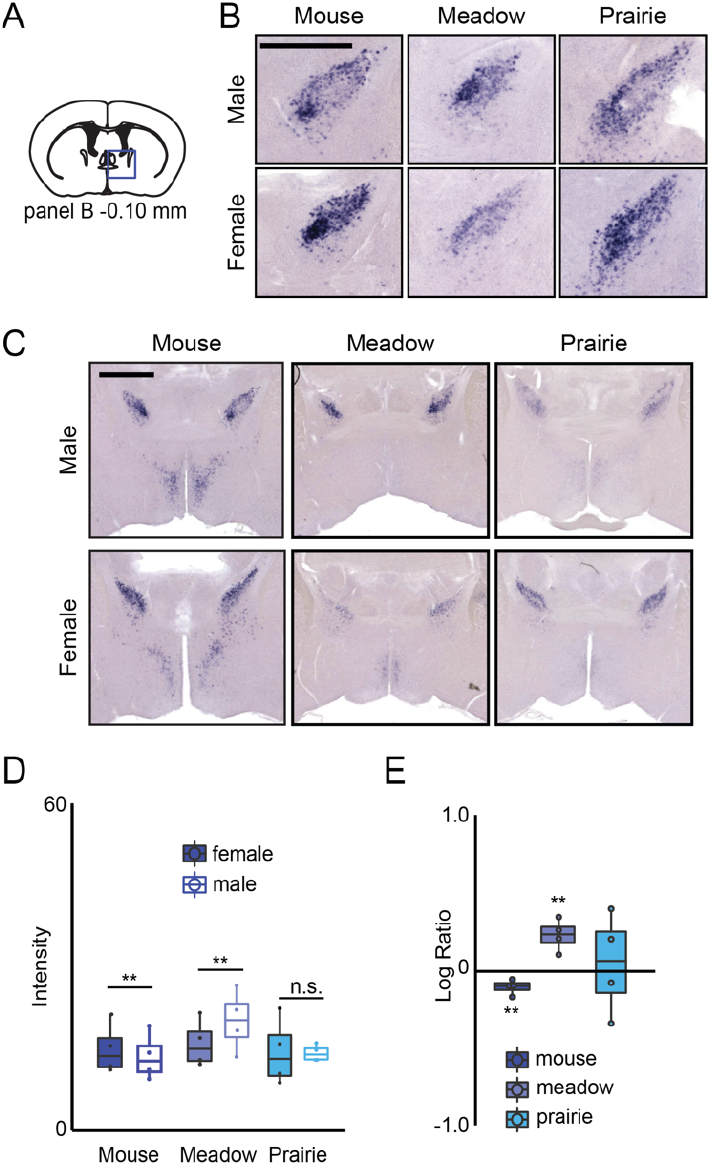
Sex differences in *Esr2* expression. **(A)** Schematic of coronal section for BNSTpr *Esr2* expression. **(B)** ISH for *Esr2* in BNSTpr. Male (top panels) and female (bottom panels) are shown for mouse, meadow vole, and prairie vole (left to right). **(C)** ISH for *Esr2* in BNST and preoptic area (POA) of the hypothalamus. **(D)** Quantification of BNSTpr *Esr2* expression in males and females of each species (N=4). Boxplot denotes median and 1st & 3rd quartiles, ** p<0.01. **(E)** Within-batch male:female log ratios of Esr2 expression in BNSTpr. ** denotes significant difference from log ratio of zero. Scale bars = 1000 μm.

### Expression of Gonadal Hormone Receptors in Mouse Hippocampus and Cortex

With the availability of hippocampal and cortical single nucleus transcriptomic data, we examined the expression of two other gonadal hormone receptor genes, progesterone receptor (*Pgr*) and G-protein coupled estrogen receptor 1 (*Gper1*), a putative membrane receptor for estrogens (Revankar et al., 2005; Urban et al., 2023), in the mouse hippocampus and cortex. We observe moderate expression of *Pgr* within the hippocampus, largely mirroring that of *Ar*, although CA2 expression was absent (Supplementary Figure 7A). In the cortex, the pattern of *Pgr* also resembles *Ar* rather than *Esr1*, with low levels in inhibitory neurons and extensive expression in excitatory populations (Supplementary Figure 7B-G). In contrast, *Gper1* was barely detectable in any neuronal types examined (Supplementary Figure 8), although we note expression is reported in SMC-Pericyte cell cluster 380 (Yao et al., 2021).

## Discussion

### Species Differences in Gonadal Hormone Receptor Expression

We find robust species differences in the distribution of gonadal hormone receptors within the hippocampus. The very low expression of *Esr1* in the mouse hippocampus corroborates previous findings from snRNA-seq data (Cembrowski et al., 2016; Yao et al., 2021) (Figure 1D) and *in situ* hybridization (Lein et al., 2007), as well as earlier characterization by radiolabeled ISH in rats (Shughrue et al., 1997; Simerly et al., 1990). One rat study reported stronger labeling in the ventral hippocampus compared to dorsal, suggesting previously unexplored species differences even within traditional laboratory models (Simerly et al., 1990). We find that prairie voles have enriched expression of *Ar* and *Esr1* within the CA3 region of the hippocampus, which is highly interconnected with the dorsal LS in mice (Besnard and Leroy, 2022). LS-HPC connectivity is important for maintaining a cognitive spatial map (Tingley and Buzsáki, 2018), therefore we suggest that variation in hippocampal *Esr1* and *Ar* expression may relate to well-documented species differences in territory usage (Getz, 1961; Madison, 1980; McGuire and Getz, 1998; Ophir et al., 2012, 2008; Streatfeild et al., 2011). Prairie voles largely share and defend a territory within a bonded pair, while meadow voles of both sexes have distinct but overlapping territories.

The LS consists entirely of GABAergic neurons and inhibits aggressive behavior via projections to the VMHvl (Albert and Chew, 1980; Wong et al., 2016). In mice, neural activity in the VMHvl is further disinhibited by excitatory input from hippocampal CA2, which can be increased by activation of vasopressin receptor 1b (Leroy et al., 2018). We find that *Ar* is expressed within the CA2 of mice, but not meadow or prairie voles, suggesting that CA2 modulation of social memory and social aggression is independent of testosterone in voles. This finding is consistent with the context-dependent selective display of aggression in both sexes of these species: meadow voles show territorial aggression during the long days of the reproductive season, while prairie voles are aggressive towards strangers only following pair bond formation (Lee and Beery, 2022). Indeed, in prairie voles, activation of the LS in pair bonded males decreases aggression towards novel conspecifics (Sailer et al., 2022). It will be interesting to determine how neural activity within CA2 to LS projections is altered in different environmental, hormonal, or social contexts.

We believe the lateral VMHvl *Esr1* population represents an expansion of the VMHvll *Cckar* neurons described in mice to be critical for female sexual behaviors (Yin et al., 2022). In prairie voles, which are induced ovulators and display less sexual dimorphism in social behavior, this neuronal population may be more responsive to estrogens in females to facilitate the expression of female sexual behaviors after puberty and after induction of ovulation by male-associated stimuli. Future studies could assess genes restricted to the medial (*Npy2r, Crhbp*) and lateral (*Cckar, Tac1*) VMHvl of mice to ascertain if the VMHvll in voles has a distinct identity (Hashikawa et al., 2017; Liu et al., 2021; Yin et al., 2022)

In general, *Ar* expression is more diffuse in voles than in mice. While the present study characterizes this expression at postnatal day 14, previous studies of circulating hormones in adults have shown that circulating testosterone is lower in group-housed prairie voles than in other vole species (Klein et al., 1997). It should be noted, however, that male prairie voles paradoxically show an increase in testosterone around the time of birth of sired litters which promotes paternal behavior, unlike in non-monogamous species (Carter and Perkeybile, 2018). This generally low testosterone at baseline and increase to promote paternal behavior may further modulate *Ar* particularly in regions where *Esr1* is coexpressed because signaling via ERα regulates the expression of the *Ar* gene (Gegenhuber et al., 2022; Hackenberg et al., 1992; Handa et al., 1996; Pelletier et al., 2004). This mechanism could also increase *Ar* expression in paired, cycling females.

However, we also see species differences beyond the overall weaker signal in voles, particularly in the hippocampus, VMH, and cortex, which suggests that the decreased expression of *Ar* could be related to species differences in social behavior and particularly pair bonding and parenting. Another rodent species, Alston’s singing mouse (*Scotinomys teguina*), expresses almost no AR protein in the hippocampus, with only a few cells in ventral CA3. Immunostaining for AR in singing mice revealed no distinct AR-immunoreactivity in the dorsal hippocampus, which contrasts with previous findings in rats and the present study in mice and voles (Simerly et al., 1990; Zheng et al., 2021). Interestingly, singing mice express more AR in the MeApv than the MeApd, contrary to the expression patterns in mice and voles described here. Although males of this species show dramatic vocal displays, which are sensitive to testosterone levels, males do not attack in a standard resident-intruder aggression paradigm (Pasch et al., 2011).

We observe no overt species differences in the pattern of *Esr2* expression, although we did not assess known sites of enrichment in the dorsal raphe and cerebellum (Nomura et al., 2005; Shughrue et al., 1997). As described previously in mice and rats, *Esr2* expression within the SBN is quite sparse and appears within a subset of *Esr1*-expressing neurons, consistent with present findings in the mouse and both vole species (Gegenhuber et al., 2022; Nomura et al., 2003; Shughrue et al., 1997). Similarly, dense expression of *Esr2* in the PVN in rats appears to be conserved in mice and voles. However, the low levels of *Esr2* reported previously in the dorsal hippocampus in rats is not seen here in mice or voles (Shughrue et al., 1997). This may be a result of the differences in methodological sensitivity between digoxigenin labeled and radiolabeled ISH hybridization; however, snRNA-seq data demonstrates virtually no *Esr2* transcripts in mouse hippocampus, and expression within only a few cortical cell types. In contrast to the low neural expression of *Esr1, Esr2*, and *Gper1, Pgr* is widespread within hippocampus and cortical excitatory populations, indicating potential for regulation of gene expression by progesterone within these brain regions.

### Decreased Sexual Divergence in Prairie Vole BNST

The distribution of the nuclear gonadal hormone receptors, *Esr1, Esr2*, and *Ar*, seems to be less sexually dimorphic in socially monogamous prairie voles than socially promiscuous meadow voles and C57BL6 mice. Species differences in the expression of these receptors may contribute to species differences in the expression of social behaviors, particularly parental behaviors, territorial aggression, and affiliative behaviors such as pair bonding. Overall, we found extensive expression of hormone receptors in the posterior/ventral BNST of both vole species. This hormone-sensitive region is critical for the coordination of social behaviors including intermale and maternal aggression, social recognition, sexual behaviors, and prosocial behaviors, and is highly interconnected with other regions of the SBN, particularly the MeA (Flanigan and Kash, 2020).

As with other mammalian species, prairie voles undergo a neonatal testosterone surge on the day of birth (Corbier et al., 1992; Lansing et al., 2013; Motelica-Heino et al., 1988), yet the impact of this surge on the brain and behavior appears minimal. We propose that precocial development in prairie voles relative to other rodents considered here may play an ontogenetic role in the decreased sexual dimorphism in this species (Wallen and Baum, 2002). Compared to mice and montane voles, prairie voles have developmentally earlier incidences of tooth eruption, eye opening, and fur growth (Shapiro and Insel, 1990). We suggest that the development of the SBN may also be precocial, leading to a partial closure of the sensitive period for brain sexual differentiation before birth. If wiring of the SBN has substantially progressed by the time neural estradiol is produced, the effects of signaling via ERα on cell survival and gene expression could be less potent, leading to less-masculinized structures and gene expression programs in the prairie vole brain. This hypothesis is consistent with the findings of BNSTpr gonadal hormone receptor expression in the present study. Less sexual dimorphism in these structures may promote selective affiliative and prosocial behaviors in males.

This hypothesis is also consistent with brain sexual differentiation in guinea pigs, the original species in which the organizational effects of testosterone were described (Phoenix et al., 1959). Although guinea pigs are also precocial, their gestation is significantly longer than that of mice, rats, or voles, ranging from 67-73 days (Connolly and Resko, 1994; Goy et al., 1964). As with rhesus macaques, the critical period for guinea pig brain sexual differentiation occurs prenatally, and is specified by fetal testosterone (Resko and Roselli, 1997). This earlier testosterone surge coincides with an earlier stage of brain development, resulting in more extensive sex differences. Our “intersectional” hypothesis predicts that the timing of developmental testosterone surges relative to brain maturation is a key driver of anatomic and behavioral sex divergence.

### Limitations and Weaknesses of the Present Study

The present study highlights a need for species-specific brain atlases for the *Microtus* genus to define species differences in anatomy, as the comparative neuroanatomy itself is poorly characterized and poses a challenge in understanding anatomical data in these species. This study is technically limited by the use of brightfield ISH; while reproducible and presenting a uniform way to study the genes of interest, brightfield ISH does not give us cellular or subcellular resolution and can only stain for one gene at a time, which prevents co-labeling to find coexpression with other markers (such as neuronal cell type markers). Furthermore, this study gives a snapshot of brain sexual differentiation across species at postnatal day 14. While we chose this developmental time point carefully based on the documented closure of the postnatal sensitive period for brain sexual differentiation in mice and rats and critical periods for social learning in rodents, exploration of perinatal expression of hormone receptors and aromatase in voles would provide additional insight into species differences in brain sexual differentiation.

## Acknowledgements

We acknowledge the CSHL Partners For the Future High School Science Research Program for support of Soyoun Moon. We thank Devanand S. Manoli (UCSF) for providing riboprobes and founder animals for our prairie vole colony, as well as Annaliese Beery (UC Berkeley/Smith) for providing founder animals for our meadow vole colony. We thank Jeremy Borniger for the use of the Keyence slide scanner and Brian Trainor for comments on the manuscript. This work was performed with assistance from CSHL LAR Shared Resource, which is funded by the Cancer Center Support Grant 5P30CA045508. We are grateful to the LAR staff who made our research with voles possible: Lisa Bianco, Jill Habel, Katherine Atehortura, Gustavo Munoz, and Rachel Rubino DVM. This work was supported by funds from the Pershing Square Foundation. This paper was typeset with the bioRxiv word template by @Chrelli: www.github.com/chrelli/bioRxiv-word-template.

## Materials and Methods

### Animals

Prairie voles (*Microtus ochrogaster*) and meadow voles (*Microtus pennsylvanicus*) were bred in-house from wild caught lineages obtained from the labs of Devanand Manoli (UCSF) and Annaliese Beery (Smith College/UC Berkeley), respectively. All voles were maintained on a 12:12 light cycle with lights on at 7am, with bedding, nesting material (nestlet), and a PVC hiding tube, and provided food and water ad libitum. C57BL6 mice were bred in-house, maintained on a 12:12 light cycle with lights on at 3am, and provided food and water ad libitum, bedding, and nesting material. All animal procedures were performed in accordance with the Cold Spring Harbor Laboratory animal care committee’s regulations.

Vole pups of both species were removed from their parental home cages and transcardially perfused with 4% paraformaldehyde (PFA) on postnatal day 14 (P14) and tails were collected for genotyping by PCR for presence or absence of the *Sry* gene. C57BL6 mouse pups were perfused at P14 and categorized by anogenital distance. Brains were extracted, cryoprotected in 30% sucrose, and stored at -80oC until sectioning for histology.

### Histology

Frozen brains were coronally cryosectioned at 50 μm, collected in RNasefree PBS, and mounted on VWR Superfrost Plus slides. In situ hybridization (ISH) using digoxigenin labeled RNA probes against androgen receptor (*Ar*), estrogen receptor alpha (*Esr1*), or estrogen receptor beta (*Esr2*) was performed as previously described (Gegenhuber et al. 2022) (details of the probes used for these genes can be found in Supplemental Table 1). Sections were post-fixed on slides with cold 4% PFA for 20 minutes, rinsed, and treated with proteinase K (10 μg/mL, Roche) for 20 minutes at room temperature. Sections were then postfixed again for 5 minutes before treatment with acetylation buffer for 10 minutes and permeabilization in 1% Triton for 10 minutes. Slides were then rinsed and sections equilibrated in hybridization solution for 2-4 hours at room temperature. Sections were subsequently incubated for 16-20 hours at 65oC in fresh hybridization buffer containing 350 ng/mL probe and washed in 0.2X SSC buffer. Washes were followed by blocking in 10% heat-inactivated sheep serum for 1-3 hours and incubation in buffer containing sheep antidigoxigenin antibody (Roche) at 1:5000 dilution for 16-20 hours at 4oC. After 2 hours of repeated washing, slides were incubated at 37oC in a staining solution containing nitro blue tetrazolium and 5-bromo-4-chloro-3-indolyl-phosphate (Roche) for 20-24 hours. Staining was stopped with 1mM EDTA, and slides were washed, postfixed, and coverslipped.

### Microscopy

All slides were imaged under brightfield illumination at 2x and/or 10x magnification and stitched using a Keyence BZ-X800 All-in-One Fluorescence microscope and Keyence BZ-X Viewer/Analyzer software. All 50μm sections were imaged from anterior (Bregma 0.38 mm) to posterior (Bregma -2.80 mm). Processed images were analyzed by an investigator blinded to the sex of the animal and mean intensity of staining was quantified and normalized. In brief, a uniform mask was applied to all individuals for each section of the region of interest as well as a background region for each image in Adobe Photoshop 2022. Using Fiji analysis software (NIH), each image was color inverted, the background staining of each image was subtracted from the ROI, and the average pixel brightness of staining was quantified. These scores were then averaged bilaterally and summed for a total region intensity.

### Statistical Analysis

Statistical analysis was performed in Excel and all plots were made using the ggplot2 (v3.4.1) package in R v4.2.1 (https://cran.r-project.org/bin/windows/base/old/4.2.1/) (Wickham, 2011). Because intensity scores within a replicate were correlated with one another, we expressed the intensities as a ratio of male to female intensity scores for each replicate. Ratios greater than 1 indicate a male bias of the gene of interest, ratios less than 1 indicate a female bias, and ratios of 1 indicate no sex bias. We then performed a one-sample t-test within each species on the log of the ratios to determine if the log ratio was significantly different from 0.

### Single Nucleus RNA Sequencing Analysis

snRNA-seq data containing 1,228,636 single-cell 10xv2 transcriptomes across several brain regions and sex and corresponding metadata were accessed from The Allen Institute for Brain Science Brain Map Portal: https://portal.brain-map.org/atlases-and-data/rnaseq/mouse-whole-cortex-and-hippocampus-10x.

To analyze expression levels of hormone receptors (*Ar, Esr1, Esr2, Pgr*, and G-protein coupled estrogen receptor 1, *Gper1*), single-cell transcriptomes were subsampled by neighborhoods and subclasses according to the cell-type taxonomy established in Yao et al., 2021. Neighborhoods (“ “) and clusters (*n_start_-n_end_*) for each subsample are as follows: 1) Hippocampus, glutamatergic: “DG/SUB/CA” 318-364; 2) Cortex, GABAergic: “CGE, MGE” 5-123; 3) Cortex, layer 2/3 glutamatergic: “L2/3 IT” 124-177; 4) Cortex, layer 4/5/6 glutamatergic: “L4/5/6 IT Car3” 178-238; 5) Cortex PT glutamatergic: “PT” 239-263; 6) Cortex layer 6 glutamatergic: “NP/CT/L6b” 264-317. Cluster labels and abbreviations used in this analysis are provided in Supplemental Table 2. Subsample gene expression counts and metadata were read into R via HDF5 (rhdf5:(Fischer et al., 2023); HDF5Array:(Pagès, 2023)) and then loaded into a Seurat object (Satija et al., 2015). Counts were LogNormalized and scaled within each subsample. Subsamples were visualized with VlnPlot() by cluster labels, then exported and designed with Adobe Illustrator.

**Supplementary Figure 1.**
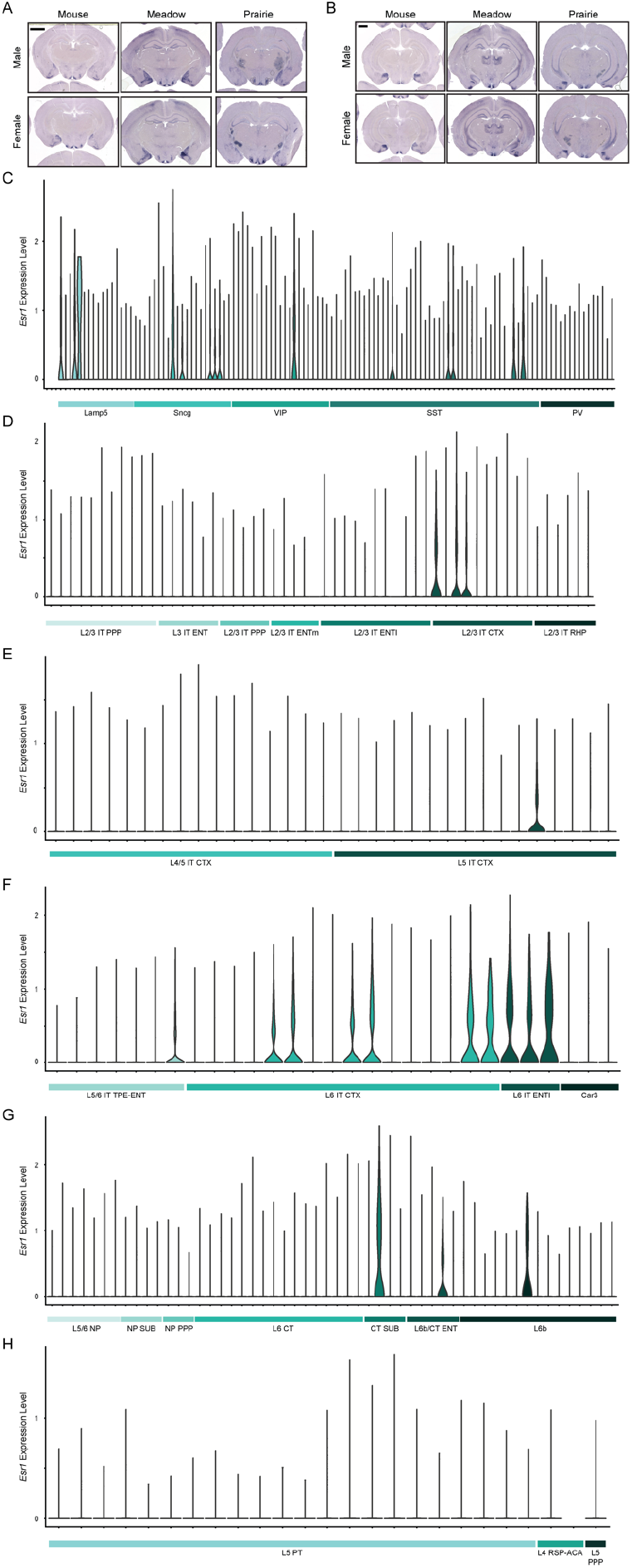
*Esr1* is sparsely expressed in hippocampus and cortex. **(A-B)** Full section views of dHPC **(A)** and vHPC **(B)** staining. Scale bars = 1000 μm. **(C-H)** Violin plots of cortical expression of *Esr1* in snRNA-seq clusters, subsampled by transcriptomic taxonomy neighborhoods (Yao et al., 2021). See Supplementary Table 2 for subclass abbreviations and cluster identity.

**Supplementary Figure 2.**
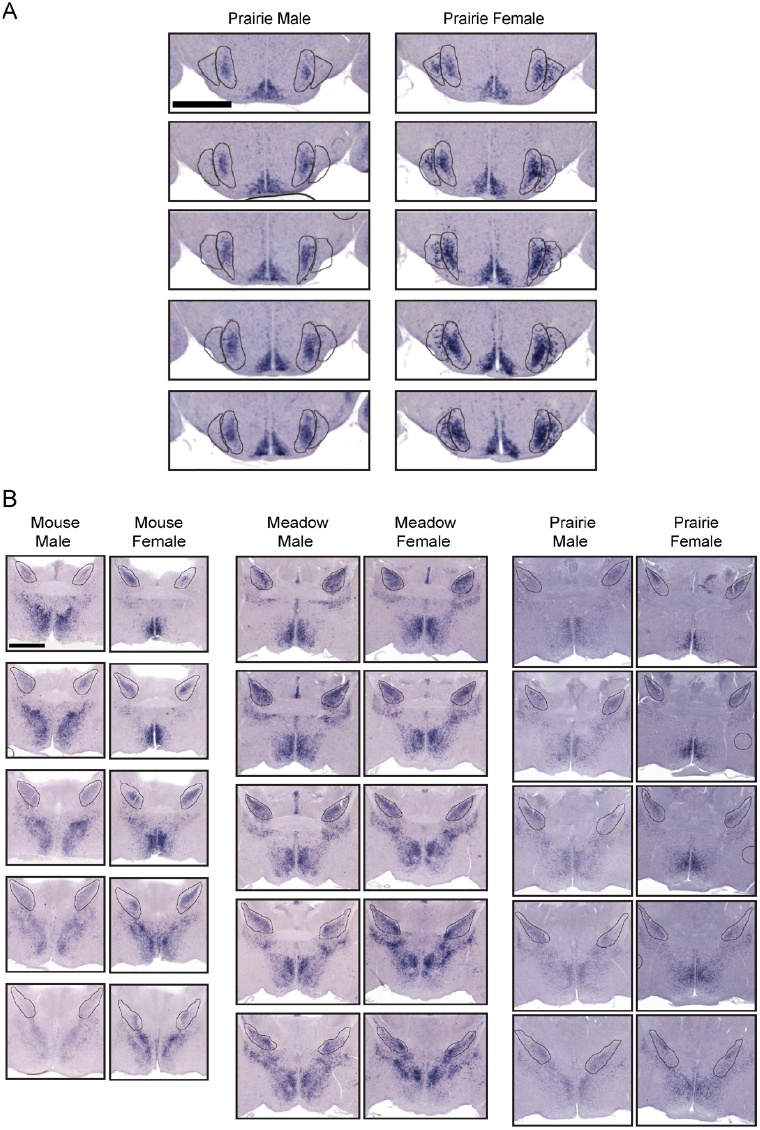
*Esr1* quantification. **(A)** Five consecutive VMHvl sections from one representative replicate for each sex of prairie voles. The VMHvl and the female-specific VMHvll region are outlined in black. **(B)** Five consecutive BNST/POA sections from one representative replicate for each sex and species. The quantified BNSTpr region is outlined in black. All BNSTpr images are to the same scale (scale bar = 1000 μm). Note the species differences in brain volume.

**Supplementary Figure S3.**
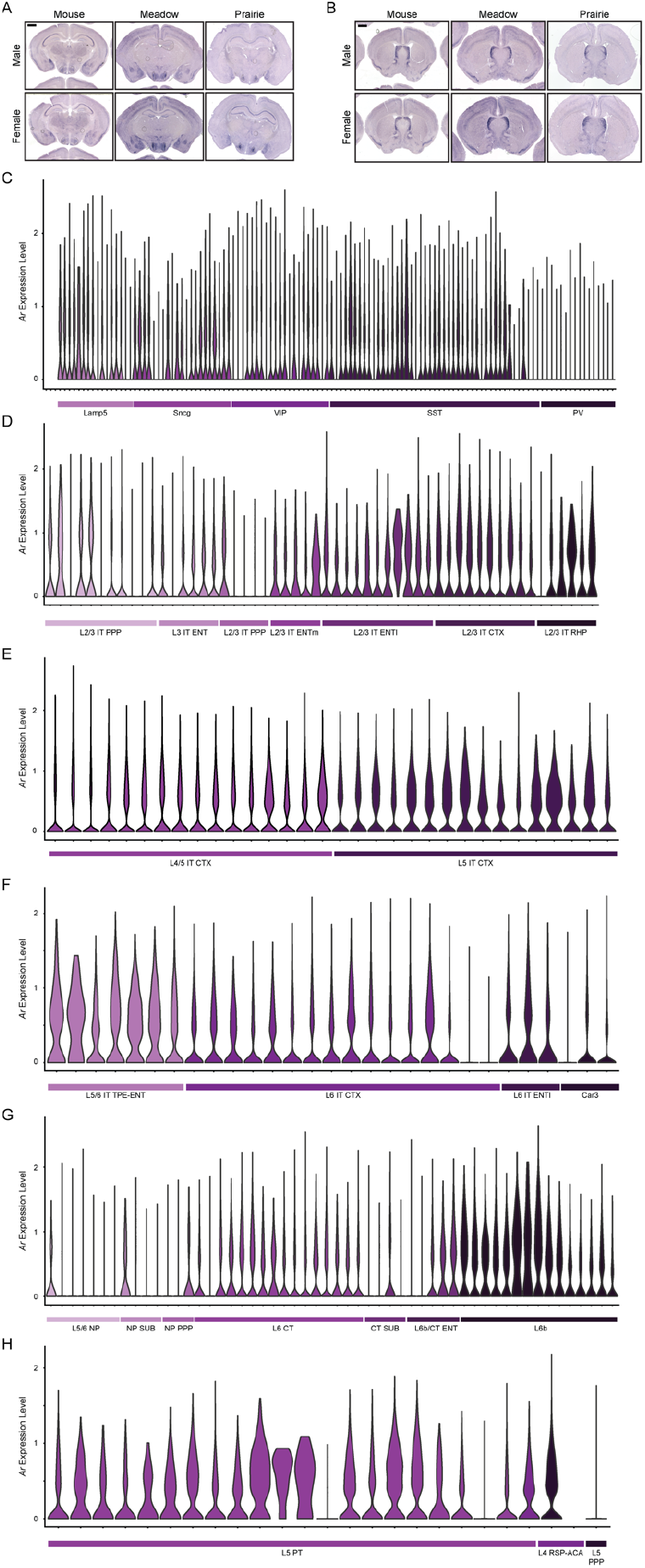
*Ar* is widely expressed in hippocampus and cortex. **(A-B)** Full section views of dHPC (A) and ACC and LS (B) staining. Scale bars = 1000 μm. **(C-H)** Violin plots of cortical expression of *Ar* in snRNA-seq clusters, subsampled by transcriptomic taxonomy neighborhoods (Yao et al., 2021). See Supplementary Table 2 for subclass abbreviations and cluster identity.

**Supplementary Figure S4.**
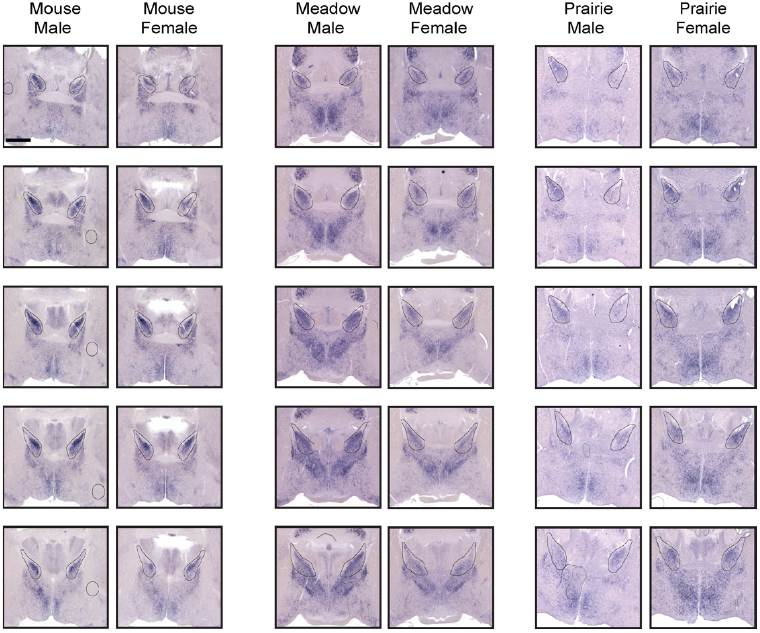
*Ar* quantification. Five consecutive BNST/POA sections from one representative replicate for each sex and species. The quantified BNSTpr region is outlined in black. All BNSTpr images are to the same scale (scale bar = 1000 μm). Note the species differences in brain volume.

**Supplementary Figure 5.**
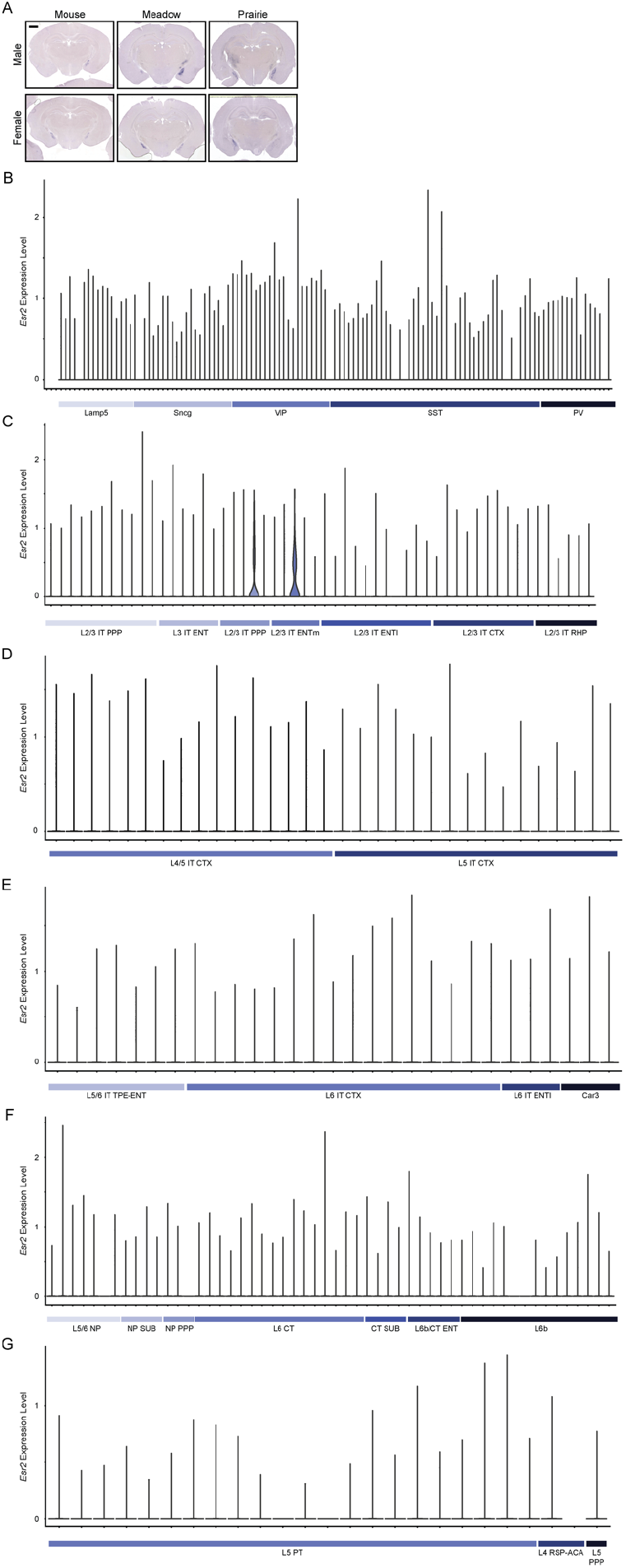
*Esr2* is sparsely expressed in hippocampus and cortex. **(A)** Full coronal sections including MeA staining. Scale bars = 1000 μm. **(B-G)** Violin plots of cortical expression of *Esr2* in snRNA-seq clusters, subsampled by transcriptomic taxonomy neighborhoods (Yao et al., 2021). See Supplementary Table 2 for subclass abbreviations and cluster identity.

**Supplementary Figure 6.**
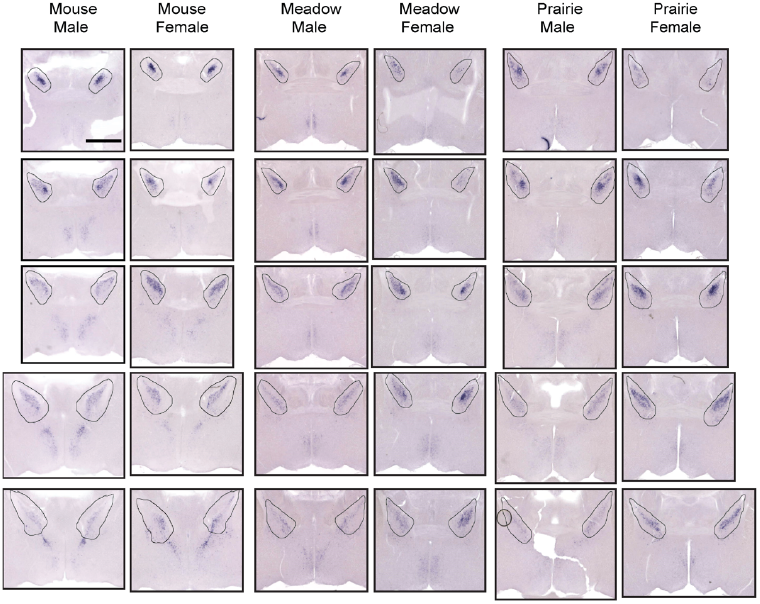
*Esr2* quantification. Five consecutive BNST/POA sections from one representative individual for each sex and species. The quantified BNSTpr region is outlined in black. Scale bars = 1000 μm. Note the species differences in brain volume.

**Supplementary Figure 7.**
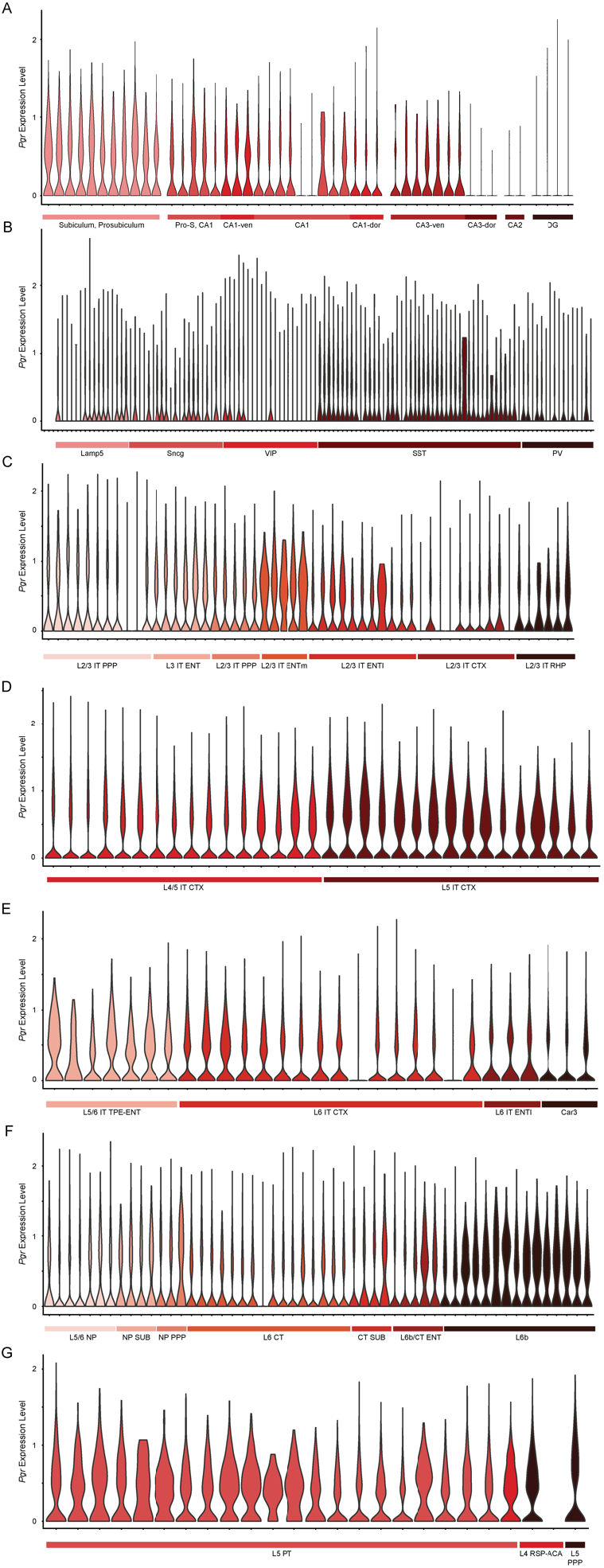
*Pgr* is widely expressed in mouse cortex and hippocampus. Violin plots across hippocampal **(A)** and cortical inhibitory (B)and excitatory **(C-H)** transcriptional cell type clusters organized by anatomical subregion (Yao et al., 2021). See Supplementary Table 2 for subclass abbreviations and cluster identity.

**Supplementary Figure 8.**
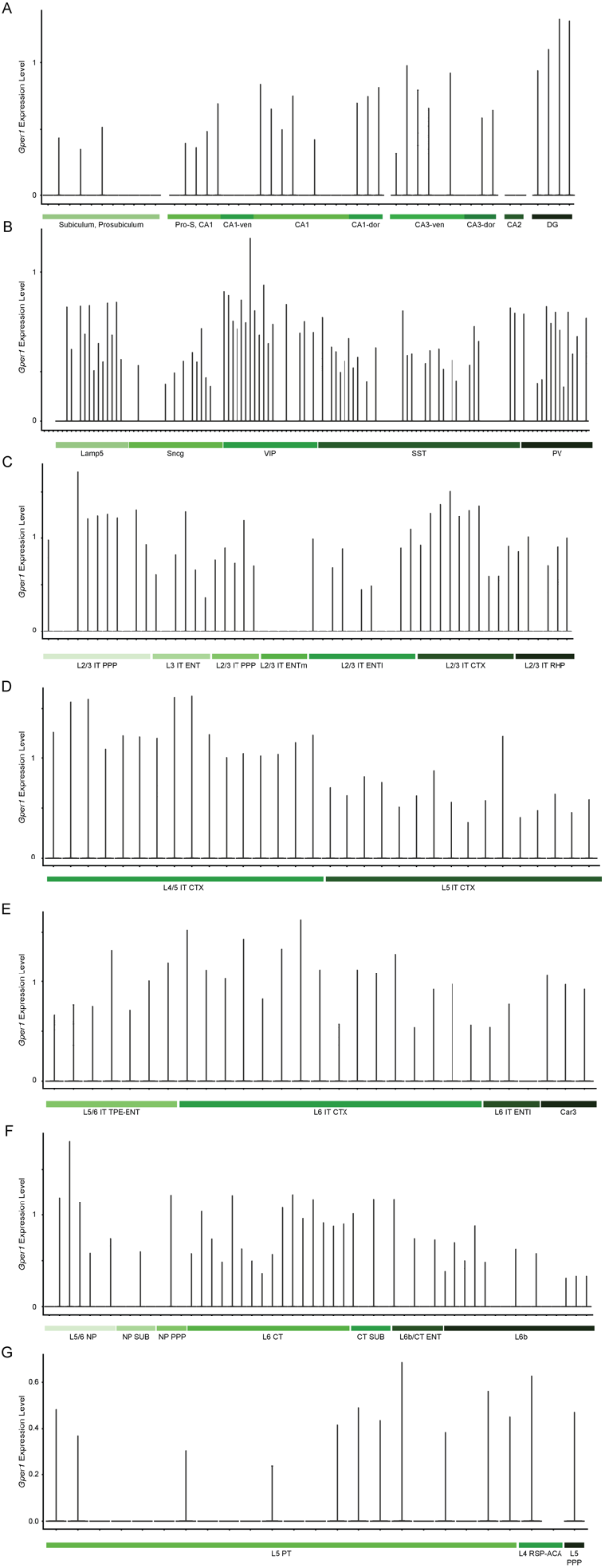
*Gper1* is sparsely expressed in mouse cortex and hippocampus. Violin plots across hippocampal **(A)** and cortical inhibitory **(B)** and excitatory **(C-H)** transcriptional cell type clusters organized by anatomical subregion (Yao et al., 2021). See Supplementary Table 2 for subclass abbreviations and cluster identity.

**Supplementary Table 1.** In situ hybridization riboprobes.

| Gene | RefSeq (vole) | Range (vole) | RefSeq (mouse) | Range (mouse) |
| --- | --- | --- | --- | --- |
| Ar | XM_013350520.2 | 2606-3285 | NM_013476.4 | 1710-2749 |
| Esr1 | NM_001282251.1 | 558-1219 | NM_007956.5 | 1152-2132 |
| Esr2 | XM_005343221.3 | 646-1309 | NM_010157.3 | 1022-1923 |

**Supplementary Table 2. Description of snRNA-seq clusters from Yao et al., 2021**.

## References

Adkins-Regan, E., 2012. Hormonal organization and activation: evolutionary implications and questions. Gen. Comp. Endocrinol. 176, 279–285.

Albert, D.J., Chew, G.L., 1980. The septal forebrain and the inhibitory modulation of attack and defense in the rat. A review. Behav. Neural Biol. 30, 357–388.

Allen, L.S., Gorski, R.A., 1990. Sex difference in the bed nucleus of the stria terminalis of the human brain. J. Comp. Neurol. 302, 697–706.

Anacker, A.M.J., Reitz, K.M., Goodwin, N.L., Beery, A.K., 2016. Stress impairs new but not established relationships in seasonally social voles. Horm. Behav. 79, 52–57.

Andersson, S., Sundberg, M., Pristovsek, N., Ibrahim, A., Jonsson, P., Katona, B., Clausson, C.-M., Zieba, A., Ramström, M., Söderberg, O., Williams, C., Asplund, A., 2017. Insufficient antibody validation challenges oestrogen receptor beta research. Nat. Commun. 8, 15840.

Bales, K.L., Kramer, K.M., Lewis-Reese, A.D., Carter, C.S., 2006. Effects of stress on parental care are sexually dimorphic in prairie voles. Physiol. Behav. 87, 424–429.

Bales, K.L., Saltzman, W., 2016. Fathering in rodents: Neurobiological substrates and consequences for offspring. Horm. Behav. 77, 249– 259.

Balthazart, J., Ball, G.F., 1998. New insights into the regulation and function of brain estrogen synthase (aromatase). Trends Neurosci. 21, 243– 249.

Beery, A.K., 2019. Frank Beach award winner: Neuroendocrinology of group living. Horm. Behav. 107, 67–75.

Beery, A.K., Vahaba, D.M., Grunberg, D.M., 2014. Corticotropin-releasing factor receptor densities vary with photoperiod and sociality. Horm. Behav. 66, 779–786.

Besnard, A., Leroy, F., 2022. Top-down regulation of motivated behaviors via lateral septum sub-circuits. Mol. Psychiatry 27, 3119–3128.

Brock, O., De Mees, C., Bakker, J., 2015. Hypothalamic expression of oestrogen receptor α and androgen receptor is sex-, age- and region-dependent in mice. J. Neuroendocrinol. 27, 264–276.

Campi, K.L., Jameson, C.E., Trainor, B.C., 2013. Sexual Dimorphism in the Brain of the Monogamous California Mouse (Peromyscus californicus). Brain Behav. Evol. 81, 236–249.

Carter, C.S., Devries, C.A., Getz, L.L., 1995. Physiological substrates of mammalian monogamy: The prairie vole model. Neurosci. Biobehav. Rev. 19, 303–314.

Carter, C.S., Getz, L.L., 1993. Monogamy and the prairie vole. Sci. Am. 268, 100–106.

Carter, C.S., Perkeybile, A.M., 2018. The Monogamy Paradox: What Do Love and Sex Have to Do With It? Front Ecol Evol 6. https://doi.org/10.3389/fevo.2018.00202

Cembrowski, M.S., Wang, L., Sugino, K., Shields, B.C., Spruston, N., 2016. Hipposeq: a comprehensive RNA-seq database of gene expression in hippocampal principal neurons. Elife 5, e14997.

Chen, P., Hong, W., 2018. Neural Circuit Mechanisms of Social Behavior. Neuron 98, 16–30.

Connolly, P.B., Resko, J.A., 1994. Prenatal testosterone differentiates brain regions controlling gonadotropin release in guinea pigs. Biol. Reprod. 51, 125–130.

Corbier, P., Edwards, D.A., Roffi, J., 1992. The neonatal testosterone surge: a comparative study. Arch. Int. Physiol. Biochim. Biophys. 100, 127– 131.

Cushing, B.S., 2016. Estrogen Receptor Alpha Distribution and Expression in the Social Neural Network of Monogamous and Polygynous Peromyscus. PLoS One 11, e0150373.

Cushing, B.S., Razzoli, M., Murphy, A.Z., Epperson, P.M., Le, W.W., Hoffman, G.E., 2004. Intraspecific variation in estrogen receptor alpha and the expression of male sociosexual behavior in two populations of prairie voles. Brain Res. 1016, 247–254.

Cushing, B.S., Wynne-Edwards, K.E., 2006. Estrogen receptor-alpha distribution in male rodents is associated with social organization. J. Comp. Neurol. 494, 595–605.

Dewsbury, D.A., 1985. Paternal behavior in rodents. Am. Zool. 25, 841–852.

Dewsbury, D.A., Baumgerdner, D.J., Evans, R.L., Webster, D.G., 1980. Sexual dimorphism for body mass in 13 taxa of muroid rodents under laboratory conditions. J. Mammal. 61, 146–149.

Fischer, B., Smith, M., Pau, G., 2023. rhdf5: R Interface to HDF5. https://doi.org/10.18129/B9.bioc.rhdf5

Flanigan, M.E., Kash, T.L., 2020. Coordination of social behaviors by the bed nucleus of the stria terminalis. Eur. J. Neurosci. https://doi.org/10.1111/ejn.14991

Gegenhuber, B., Wu, M.V., Bronstein, R., Tollkuhn, J., 2022. Gene regulation by gonadal hormone receptors underlies brain sex differences. Nature 1–7.

Getz, L.L., 1961. Home Ranges, Territoriality, and Movement of the Meadow Vole. J. Mammal. 42, 24–36.

Getz, L.L., Carter, C.S., Gavish, L., 1981. The mating system of the prairie vole, Microtus ochrogaster: Field and laboratory evidence for pairbonding. Behav. Ecol. Sociobiol. 8, 189–194.

Goodson, J.L., 2005. The vertebrate social behavior network: evolutionary themes and variations. Horm. Behav. 48, 11–22.

Gorski, R.A., Gordon, J.H., Shryne, J.E., Southam, A.M., 1978. Evidence for a morphological sex difference within the medial preoptic area of the rat brain. Brain Res. 148, 333–346.

Goy, R.W., Bridson, W.E., Young, W.C., 1964. PERIOD OF MAXIMAL SUSCEPTIBILITY OF THE PRENATAL FEMALE GUINEA PIG TO MASCULINIZING ACTIONS OF TESTOSTERONE PROPIONATE. J. Comp. Physiol. Psychol. 57, 166–174.

Hackenberg, R., Hawighorst, T., Filmer, A., Slater, E.P., Bock, K., Beato, M., Schulz, K.D., 1992. Regulation of androgen receptor mRNA and protein level by steroid hormones in human mammary cancer cells. J. Steroid Biochem. Mol. Biol. 43, 599–607.

Hammock, E.A., 2014. Developmental Perspectives on Oxytocin and Vasopressin. Neuropsychopharmacology 40, 24–42.

Handa, R.J., Kerr, J.E., DonCarlos, L.L., McGivern, R.F., Hejna, G., 1996. Hormonal regulation of androgen receptor messenger RNA in the medial preoptic area of the male rat. Brain Res. Mol. Brain Res. 39, 57– 67.

Hashikawa, K., Hashikawa, Y., Lischinsky, J., Lin, D., 2018. The Neural Mechanisms of Sexually Dimorphic Aggressive Behaviors. Trends Genet. 34, 755–776.

Hashikawa, K., Hashikawa, Y., Tremblay, R., Zhang, J., Feng, J.E., Sabol, A., Piper, W.T., Lee, H., Rudy, B., Lin, D., 2017. Esr1+ cells in the ventromedial hypothalamus control female aggression. Nat. Neurosci. 20, 1580–1590.

He, Z., Young, L., Ma, X.-M., Guo, Q., Wang, L., Yang, Y., Luo, L., Yuan, W., Li, L., Zhang, J., Hou, W., Qiao, H., Jia, R., Tai, F., 2019. Increased anxiety and decreased sociability induced by paternal deprivation involve the PVN-PrL OTergic pathway. Elife 8. https://doi.org/10.7554/eLife.44026

Heske, E.J., Ostfeld, R.S., 1990. Sexual Dimorphism in Size, Relative Size of Testes, and Mating Systems in North American Voles. J. Mammal. 71, 510–519.

Hnatczuk, O.C., Lisciotto, C. A., DonCarlos, L.L., Carter, C.S., Morrell, J.I., 1994. Estrogen receptor immunoreactivity in specific brain areas of the prairie vole (Microtus ochrogaster) is altered by sexual receptivity and genetic sex. J. Neuroendocrinol. 6, 89–100.

Hoke, K.L., Adkins-Regan, E., Bass, A.H., McCune, A.R., Wolfner, M.F., 2019. Co-opting evo-devo concepts for new insights into mechanisms of behavioural diversity. J. Exp. Biol. 222. https://doi.org/10.1242/jeb.190058

Ikeda, Y., Nagai, A., Ikeda, M.-A., Hayashi, S., 2003. Sexually dimorphic and estrogen-dependent expression of estrogen receptor beta in the ventromedial hypothalamus during rat postnatal development. Endocrinology 144, 5098–5104.

Insel, T.R., Gelhard, R., Shapiro, L.E., 1991. The comparative distribution of forebrain receptors for neurohypophyseal peptides in monogamous and polygamous mice. Neuroscience 43, 623–630.

Insel, T.R., Young, L.J., 2001. The neurobiology of attachment. Nat. Rev. Neurosci. 2, 129–136.

Juntti, S.A., Tollkuhn, J., Wu, M.V., Fraser, E.J., Soderborg, T., Tan, S., Honda, S.-I., Harada, N., Shah, N.M., 2010. The androgen receptor governs the execution, but not programming, of male sexual and territorial behaviors. Neuron 66, 260–272.

Kelly, A.M., Vitousek, M.N., 2017. Dynamic modulation of sociality and aggression: an examination of plasticity within endocrine and neuroendocrine systems. Philos. Trans. R. Soc. Lond. B Biol. Sci. 372. https://doi.org/10.1098/rstb.2016.0243

Kelly, D.A., Varnum, M.M., Krentzel, A.A., Krug, S., Forger, N.G., 2013. Differential control of sex differences in estrogen receptor α in the bed nucleus of the stria terminalis and anteroventral periventricular nucleus. Endocrinology 154, 3836–3846.

Klein, S.L., Hairston, J.E., Devries, A.C., Nelson, R.J., 1997. Social environment and steroid hormones affect species and sex differences in immune function among voles. Horm. Behav. 32, 30–39.

Kramer, K.M., Perry, A.N., Golbin, D., Cushing, B.S., 2009. Sex steroids are necessary in the second postnatal week for the expression of male alloparental behavior in prairie voles (Microtus ochragaster). Behav. Neurosci. 123, 958–963.

Lansing, S.W., French, J.A., Lonstein, J.S., 2013. Circulating plasma testosterone during early neonatal life in the socially monogamous and biparental prairie vole (Microtus ochrogaster). Psychoneuroendocrinology 38, 306–309.

Lee, N.S., Beery, A.K., 2022. Selectivity and Sociality: Aggression and Affiliation Shape Vole Social Relationships. Front. Behav. Neurosci. 16, 826831.

Lee, N.S., Beery, A.K., 2019. Neural Circuits Underlying Rodent Sociality: A Comparative Approach. Curr. Top. Behav. Neurosci. 43, 211–238.

Lein, E.S., Hawrylycz, M.J., Ao, N., Ayres, M., Bensinger, A., Bernard, A., Boe, A.F., Boguski, M.S., Brockway, K.S., Byrnes, E.J., Chen, Lin, Chen, Li, Chen, T.-M., Chin, M.C., Chong, J., Crook, B.E., Czaplinska, A., Dang, C.N., Datta, S., Dee, N.R., Desaki, A.L., Desta, T., Diep, E., Dolbeare, T.A., Donelan, M.J., Dong, H.-W., Dougherty, J.G., Duncan, B.J., Ebbert, A.J., Eichele, G., Estin, L.K., Faber, C., Facer, B.A., Fields, R., Fischer, S.R., Fliss, T.P., Frensley, C., Gates, S.N., Glattfelder, K.J., Halverson, K.R., Hart, M.R., Hohmann, J.G., Howell, M.P., Jeung, D.P., Johnson, R.A., Karr, P.T., Kawal, R., Kidney, J.M., Knapik, R.H., Kuan, C.L., Lake, J.H., Laramee, A.R., Larsen, K.D., Lau, C., Lemon, T.A., Liang, A.J., Liu, Y., Luong, L.T., Michaels, J., Morgan, J.J., Morgan, R.J., Mortrud, M.T., Mosqueda, N.F., Ng, L.L., Ng, R., Orta, G.J., Overly, C.C., Pak, T.H., Parry, S.E., Pathak, S.D., Pearson, O.C., Puchalski, R.B., Riley, Z.L., Rockett, H.R., Rowland, S.A., Royall, J.J., Ruiz, M.J., Sarno, N.R., Schaffnit, K., Shapovalova, N.V., Sivisay, T., Slaughterbeck, C.R., Smith, S.C., Smith, K.A., Smith, B.I., Sodt, A.J., Stewart, N.N., Stumpf, K.-R., Sunkin, S.M., Sutram, M., Tam, A., Teemer, C.D., Thaller, C., Thompson, C.L., Varnam, L.R.,Visel, A., Whitlock, R.M., Wohnoutka, P.E., Wolkey, C.K., Wong, V.Y., Wood, M., Yaylaoglu, M.B., Young, R.C., Youngstrom, B.L., Yuan, X.F., Zhang, B., Zwingman, T.A., Jones, A.R., 2007. Genome-wide atlas of gene expression in the adult mouse brain. Nature 445, 168–176.

Lephart, E.D., 1996. A review of brain aromatase cytochrome P450. Brain Res. Brain Res. Rev. 22, 1–26.

Leroy, F., Park, J., Asok, A., Brann, D.H., Meira, T., Boyle, L.M., Buss, E.W., Kandel, E.R., Siegelbaum, S.A., 2018. A circuit from hippocampal CA2 to lateral septum disinhibits social aggression. Nature 564, 213–218.

Lim, M.M., Wang, Z., Olazábal, D.E., Ren, X., Terwilliger, E.F., Young, L.J., 2004. Enhanced partner preference in a promiscuous species by manipulating the expression of a single gene. Nature 429, 754–757.

Liu, M., Kim, D.-W., Zeng, H., Anderson, D.J., 2021. Make war not love: The neural substrate underlying a state-dependent switch in female social behavior. Neuron. https://doi.org/10.1016/j.neuron.2021.12.002

Lonstein, J.S., De Vries, G.J., 2000. Sex differences in the parental behavior of rodents. Neurosci. Biobehav. Rev. 24, 669–686.

Lonstein, J.S., Gammie, S.C., 2002. Sensory, hormonal, and neural control of maternal aggression in laboratory rodents. Neurosci. Biobehav. Rev. 26, 869–888.

Lonstein, J.S., Rood, B.D., De Vries, G.J., 2002. Parental responsiveness is feminized after neonatal castration in virgin male prairie voles, but is not masculinized by perinatal testosterone in virgin females. Horm. Behav. 41, 80–87.

MacLusky, N.J., Naftolin, F., 1981. Sexual differentiation of the central nervous system. Science 211, 1294–1302.

Madison, D.M., 1980. Space use and social structure in meadow voles, Microtus pennsylvanicus. Behav. Ecol. Sociobiol. 7, 65–71.

McCarthy, M.M., 2008. Estradiol and the developing brain. Physiol. Rev. 88, 91–124.

McGuire, B., Getz, L.L., 1998. The nature and frequency of social interactions among free-living prairie voles (Microtus ochrogaster). Behav. Ecol. Sociobiol. 43, 271–279.

Motelica-Heino, I., Castanier, M., Corbier, P., Edwards, D.A., Roffi, J., 1988. Testosterone levels in plasma and testes of neonatal mice. J. Steroid Biochem. 31, 283–286.

Naftolin, F., Ryan, K.J., 1975. The metabolism of androgens in central neuroendocrine tissues. J. Steroid Biochem. 6, 993–997.

Nelson, A.W., Groen, A.J., Miller, J.L., Warren, A.Y., Holmes, K.A., Tarulli, G.A., Tilley, W.D., Katzenellenbogen, B.S., Hawse, J.R., Gnanapragasam, V.J., Carroll, J.S., 2017. Comprehensive assessment of estrogen receptor beta antibodies in cancer cell line models and tissue reveals critical limitations in reagent specificity. Mol. Cell. Endocrinol. 440, 138–150.

Newman, S.W., 1999. The medial extended amygdala in male reproductive behavior. A node in the mammalian social behavior network. Ann. N. Y. Acad. Sci. 877, 242–257.

Newmaster, K.T., Nolan, Z.T., Chon, U., Vanselow, D.J., Weit, A.R., Tabbaa, M., Hidema, S., Nishimori, K., Hammock, E.A.D., Kim, Y., 2020. Quantitative cellular-resolution map of the oxytocin receptor in postnatally developing mouse brains. Nat. Commun. 11, 1885.

Nomura, M., Akama, K.T., Alves, S.E., Korach, K.S., Gustafsson, J.-A., Pfaff, D.W., Ogawa, S., 2005. Differential distribution of estrogen receptor (ER)-alpha and ER-beta in the midbrain raphe nuclei and periaqueductal gray in male mouse: Predominant role of ER-beta in midbrain serotonergic systems. Neuroscience 130, 445–456.

Nomura, M., Korach, K.S., Pfaff, D.W., Ogawa, S., 2003. Estrogen receptor beta (ERbeta) protein levels in neurons depend on estrogen receptor alpha (ERalpha) gene expression and on its ligand in a brain regionspecific manner. Brain Res. Mol. Brain Res. 110, 7–14.

Oliveras, D., Novak, M., 1986. A comparison of paternal behaviour in the meadow vole Microtus pennsylvanicus, the pine vole M. pinetorum and the prairie vole M. ochrogaster. Anim. Behav. 34, 519–526.

Ophir, A.G., Gessel, A., Zheng, D.-J., Phelps, S.M., 2012. Oxytocin receptor density is associated with male mating tactics and social monogamy. Horm. Behav. 61, 445–453.

Ophir, A.G., Wolff, J.O., Phelps, S.M., 2008. Variation in neural V1aR predicts sexual fidelity and space use among male prairie voles in semi-natural settings. Proc. Natl. Acad. Sci. U. S. A. 105, 1249–1254.

Pagès, H., 2023. HDF5Array: HDF5 backend for DelayedArray objects. https://doi.org/10.18129/B9.bioc.HDF5Array

Pasch, B., George, A.S., Hamlin, H.J., Guillette, L.J., Jr, Phelps, S.M., 2011. Androgens modulate song effort and aggression in Neotropical singing mice. Horm. Behav. 59, 90–97.

Paxinos, G., Franklin, K.B.J., 2019. Paxinos and Franklin’s the Mouse Brain in Stereotaxic Coordinates. Academic Press.

Pelletier, G., Luu-The, V., Li, S., Labrie, F., 2004. Localization and estrogenic regulation of androgen receptor mRNA expression in the mouse uterus and vagina. J. Endocrinol. 180, 77–85.

Phoenix, C.H., Goy, R.W., Gerall, A.A., Young, W.C., 1959. Organizing action of prenatally administered testosterone propionate on the tissues mediating mating behavior in the female guinea pig. Endocrinology 504, 369–382.

Quadros, P.S., Wagner, C.K., 2008. Regulation of progesterone receptor expression by estradiol is dependent on age, sex and region in the rat brain. Endocrinology 149, 3054–3061.

Resko, J.A., Roselli, C.E., 1997. Prenatal hormones organize sex differences of the neuroendocrine reproductive system: observations on guinea pigs and nonhuman primates. Cell. Mol. Neurobiol. 17, 627–648.

Revankar, C.M., Cimino, D.F., Sklar, L.A., Arterburn, J.B., Prossnitz, E.R., 2005. A transmembrane intracellular estrogen receptor mediates rapid cell signaling. Science 307, 1625–1630.

Roberts, R.L., Zullo, A.S., Carter, C.S., 1997. Sexual differentiation in prairie voles: the effects of corticosterone and testosterone. Physiol. Behav. 62, 1379–1383.

Sadino, J.M., Donaldson, Z.R., 2018. Prairie Voles as a Model for Understanding the Genetic and Epigenetic Regulation of Attachment Behaviors. ACS Chem. Neurosci. 9, 1939–1950.

Sailer, L.L., Park, A.H., Galvez, A., Ophir, A.G., 2022. Lateral septum DREADD activation alters male prairie vole prosocial and antisocial behaviors, not partner preferences. Commun Biol 5, 1299.

Satija, R., Farrell, J.A., Gennert, D., Schier, A.F., Regev, A., 2015. Spatial reconstruction of single-cell gene expression data. Nat. Biotechnol. 33, 495–502.

Shapiro, L.E., Insel, T.R., 1990. Infant’s response to social separation reflects adult differences in affiliative behavior: A comparative developmental study in prairie and montane voles. Dev. Psychobiol. 23, 375–393.

Shapiro, L.E., Leonard, C.M., Sessions, C.E., Dewsbury, D.A., Insel, T.R., 1991. Comparative neuroanatomy of the sexually dimorphic hypothalamus in monogamous and polygamous voles. Brain Res. 541, 232–240.

Shughrue, P.J., Lane, M.V., Merchenthaler, I., 1997. Comparative distribution of estrogen receptor-alpha and -beta mRNA in the rat central nervous system. J. Comp. Neurol. 388, 507–525.

Simerly, R.B., 2002. Wired for reproduction: organization and development of sexually dimorphic circuits in the mammalian forebrain. Annu. Rev. Neurosci. 25, 507–536.

Simerly, R.B., Chang, C., Muramatsu, M., Swanson, L.W., 1990. Distribution of androgen and estrogen receptor mRNA-containing cells in the rat brain: an in situ hybridization study. J. Cop. Neurol. 294, 76–95.

Simerly, R.B., Swanson, L.W., Handa, R.J., Gorski, R.A., 1985. Influence of perinatal androgen on the sexually dimorphic distribution of tyrosine hydroxylase-immunoreactive cells and fibers in the anteroventral periventricular nucleus of the rat. Neuroendocrinology 40, 501–510.

St. John, R.D., Corning P.A., 1973. Maternal aggression in mice. Behav. Biol. 9, 635–639.

Streatfeild, C.A., Mabry, K.E., Keane, B., Crist, T.O., Solomon, N.G., 2011. Intraspecific variability in the social and genetic mating systems of prairie voles, Microtus ochrogaster. Anim. Behav. 82, 1387–1398.

Tickerhoof, M.C., Hale, L.H., Butler, M.J., Smith, A.S., 2020. Regulation of defeat-induced social avoidance by medial amygdala DRD1 in male and female prairie voles. Psychoneuroendocrinology 113, 104542.

Tingley, D., Buzsáki, G., 2018. Transformation of a Spatial Map across the Hippocampal-Lateral Septal Circuit. Neuron 98, 1229–1242.e5.

Tsukahara, S., Morishita, M., 2020. Sexually Dimorphic Formation of the Preoptic Area and the Bed Nucleus of the Stria Terminalis by Neuroestrogens. Front. Neurosci. 14, 797.

Urban, N., Leonhardt, M., Schaefer, M., 2023. Multiplex G Protein-Coupled Receptor Screen Reveals Reliably Acting Agonists and a Gq-Phospholipase C Coupling Mode of GPR30/GPER1. Mol. Pharmacol. 103, 48–62.

Wallace, K.J., Chun, E.K., Manns, J.R., Ophir, A.G., Kelly, A.M., 2023. A test of the social behavior network reveals differential patterns of neural responses to social novelty in bonded, but not non-bonded, male prairie voles. Horm. Behav. 152, 105362.

Wallen, K., Baum, M.J., 2002. 69 - Masculinization and Defeminization in Altricial and Precocial Mammals: Comparative Aspects of Steroid Hormone Action, in: Pfaff, D.W., Arnold, A.P., Fahrbach, S.E., Etgen, A.M., Rubin, R.T. (Eds.), Hormones, Brain and Behavior. Academic Press, San Diego, pp. 385–423.

Wang, Z., Hulihan, T.J., Insel, T.R., 1997. Sexual and social experience is associated with different patterns of behavior and neural activation in male prairie voles. Brain Res. 767, 321–332.

Wang, Z., Zhou, L., Hulihan, T.J., Insel, T.R., 1996. Immunoreactivity of central vasopressin and oxytocin pathways in microtine rodents: a quantitative comparative study. J. Comp. Neurol. 366, 726–737.

Wei, D., Talwar, V., Lin, D., 2021. Neural circuits of social behaviors: Innate yet flexible. Neuron 109, 1600–1620.

Wickham, H., 2011. Ggplot2. Wiley Interdiscip. Rev. Comput. Stat. 3, 180– 185.

Williams, J.R., Insel, T.R., Harbaugh, C.R., Carter, C.S., 1994. Oxytocin administered centrally facilitates formation of a partner preference in female prairie voles (Microtus ochrogaster). J. Neuroendocrinol. 6, 247–250.

Winslow, J.T., Hastings, N., Carter, C.S., Harbaugh, C.R., Insel, T.R., 1993. A role for central vasopressin in pair bonding in monogamous prairie voles. Nature 365, 545–548.

Wong, L.C., Wang, L., D’Amour, J.A., Yumita, T., Chen, G., Yamaguchi, T., Chang, B.C., Bernstein, H., You, X., Feng, J.E., Froemke, R.C., Lin, D., 2016. Effective Modulation of Male Aggression through Lateral Septum to Medial Hypothalamus Projection. Curr. Biol. 26, 593–604.

Wu, M.V., Manoli, D.S., Fraser, E.J., Coats, J.K., Tollkuhn, J., Honda, S.-I., Harada, N., Shah, N.M., 2009. Estrogen Masculinizes Neural Pathways and Sex-Specific Behaviors. Cell 139, 61–72.

Xu, X., Coats, J.K., Yang, C.F., Wang, A., Ahmed, O.M., Alvarado, M., Izumi, T., Shah, N.M., 2012. Modular Genetic Control of Sexually Dimorphic Behaviors. Cell 148, 596–607.

Yang, C.F., Chiang, M.C., Gray, D.C., Prabhakaran, M., Alvarado, M., Juntti, S. A., Unger, E.K., Wells, J. A., Shah, N.M., 2013. Sexually dimorphic neurons in the ventromedial hypothalamus govern mating in both sexes and aggression in males. Cell 153, 896–909.

Yao, Z., van Velthoven, C.T.J., Nguyen, T.N., Goldy, J., Sedeno-Cortes, A.E., Baftizadeh, F., Bertagnolli, D., Casper, T., Chiang, M., Crichton, K., Ding, S.-L., Fong, O., Garren, E., Glandon, A., Gouwens, N.W., Gray, J., Graybuck, L.T., Hawrylycz, M.J., Hirschstein, D., Kroll, M., Lathia, K., Lee, C., Levi, B., McMillen, D., Mok, S., Pham, T., Ren, Q., Rimorin, C., Shapovalova, N., Sulc, J., Sunkin, S.M., Tieu, M., Torkelson, A., Tung, H., Ward, K., Dee, N., Smith, K.A., Tasic, B., Zeng, H., 2021. A taxonomy of transcriptomic cell types across the isocortex and hippocampal formation. Cell 184, 3222–3241.e26.

Yin, L., Hashikawa, K., Hashikawa, Y., Osakada, T., Lischinsky, J.E., Diaz, V., Lin, D., 2022. VMHvllCckar cells dynamically control female sexual behaviors over the reproductive cycle. Neuron 110, 3000–3017.e8.

Yokosuka, M., Okamura, H., Hayashi, S., 1997. Postnatal development and sex difference in neurons containing estrogen receptor-alpha immunoreactivity in the preoptic brain, the diencephalon, and the amygdala in the rat. J. Comp. Neurol. 389, 81–93.

Young, K.A., Gobrogge, K.L., Liu, Y., Wang, Z., 2011. The neurobiology of pair bonding: insights from a socially monogamous rodent. Front. Neuroendocrinol. 32, 53–69.

Young, L.J., Crews, D., 1995. Comparative neuroendocrinology of steroid receptor gene expression and regulation: Relationship to physiology and behavior. Trends Endocrinol. Metab. 6, 317–323.

Zheng, D.-J., Singh, A., Phelps, S.M., 2021. Conservation and dimorphism in androgen receptor distribution in Alston’s singing mouse (Scotinomys teguina). J. Comp. Neurol. https://doi.org/10.1002/cne.25108

Zhou, X., Li, A., Mi, X., Li, Yixuan, Ding, Z., An, M., Chen, Y., Li, W., Tao, X., Chen, X., Li, Ying, 2023. Hyperexcited limbic neurons represent sexual satiety and reduce mating motivation. Science 379, 820–825.

Zilkha, N., Scott, N., Kimchi, T., 2017. Sexual Dimorphism of Parental Care: From Genes to Behavior. Annu. Rev. Neurosci. 40, 273–305.

Zuloaga, D.G., Zuloaga, K.L., Hinds, L.R., Carbone, D.L., Handa, R.J., 2014. Estrogen receptor β expression in the mouse forebrain: age and sex differences. J. Comp. Neurol. 522, 358–371.

